# A molecular switch: Ro5-3335 drives hematopoietic stem cell division and clonality through distinct RUNX1 binding partners

**DOI:** 10.1101/2024.06.07.596542

**Authors:** Anne L. Robertson, Lu Yue, Avik Choudhuri, Daniel Fernandez-Perez, Alex Maya-Pombo, Caroline Kubaczka, Samuel J. Wattrus, Kyle D. Drake, Joseph Mandelbaum, Serine Avagyan, Song Yang, Rohan Bhattacharya, Rebecca J. Freeman, Victoria Chan, Megan C. Blair, George Q. Daley, Alejo E. Rodriguez-Fraticelli, Leonard I. Zon

## Abstract

A defined number of hematopoietic stem cell (HSC) clones are born during development and expand to form the pool of adult stem cells. An intricate balance between self-renewal and differentiation of these HSCs supports hematopoiesis for life. Increasing the diversity of stem cell clones could have great impact on the hematopoietic stress response, for instance during chemotherapy, gene therapy or transplantation. Chemical screening for modulators of HSC expansion in zebrafish led us to identify Ro5-3335, which is known to bind to the transcription factor RUNX1. Diverse RUNX1 binding partners, including CBFβ, allow specific regulation of downstream gene expression. RUNX1 is required for definitive hematopoiesis, and its loss has been shown to reduce clonal diversity. We find that Ro5-3335 expands HSCs in zebrafish by inducing more symmetric cell divisions, leading to an increase in HSC clone number. Ro5-3335 also enhances clonal diversity in mouse HSC cultures. Using human CD34+ HSPCs, our studies reveal that Ro5-3335 remodels the RUNX1 transcription complex by a novel recruitment of ELF1/2, without altering CBFβ DNA binding affinity. This results in specific expression of cell cycle and hematopoietic genes that enhance HSC self-renewal and prevent differentiation. Our work provides the first reported evidence of an increase in HSC clone number by chemical induction *in vivo*, via novel recruitment of ELF1/2 to RUNX1, which could help develop treatments that enhance clonal diversity for blood diseases.

## Introduction

The self-renewal and differentiation potency of blood stem cells maintains hematopoiesis for life. During embryogenesis, hematopoietic stem cells (HSCs) arise from the developing aorta-gonad-mesonephros (AGM) through the process of endothelial-to-hematopoietic transition (EHT)^1–3^, then migrate to an intermediate stem cell niche where they divide. Advances in genetic and color barcoding systems have made it possible to estimate that approximately 60-190 HSC clones are born in mice^4,5^, and 20-30 HSC clones in zebrafish^6^. The diversity of these embryonic clones can be altered by extrinsic and intrinsic factors, which can have profound consequences on their contribution to adult blood. For instance, clone diversity is reduced by environmental stressors such as irradiation and transplantation, which lead to clonal selection and reduced clonal diversity^6^. Using cellular barcoding, we found that macrophages within the stem cell niche have the ability to quality assure HSCs, and removing embryonic macrophages results in only half the number of stem cell clones being maintained to adulthood^7^. Macrophages sense the surface “eat-me” signal, Calr, and the “don’t eat me” signal, B2m, to either remove an HSC or allow it to proliferate^8^. These findings highlight the role of cells within the niche microenvironment in shaping the clonal composition and long-term functionality of the HSC pool.

HSC clonality is also affected by genetic mutations. Zebrafish embryos with a mutation in Runx1 generate very low numbers of hematopoietic stem and progenitor cells (HSPCs) during development^9^, and fewer clones contribute to adult blood^10^. Runx1 is a transcription factor with a critical role in HSC development^11^, and deficiency of Runx1 in mice leads to loss of all definitive hematopoietic cells^2,12–14^. Runx1 binds to DNA via an N-terminal Runt domain, which also mediates interaction with the non-DNA binding cofactor, core-binding factor-β (CBFβ)^15^. Formation of the Runx1/CBFβ heterodimeric complex increases the DNA binding affinity of Runx1^13,16^. Unlike Runx1 deficiency, loss of CBFβ does not completely block definitive hematopoiesis. Erythroid and myeloid progenitors can be found in the fetal liver of Cbfβ deficient embryos^17^ and expression of *cmyb* is still detected in *cbfb*^−/−^ mutant zebrafish embryos^18^. This suggests that low-affinity binding of Runx1 to DNA in the absence of Cbfβ is sufficient for a low level of definitive hematopoiesis and that Runx1 and Cbfβ have distinct roles in transcriptional regulation. Runx1 can drive early emergence of HSCs even in the absence of Cbfβ, but Cbfβ is required for their release into the circulation, leading to accumulation of HSCs in the AGM in *cbfb*^−/−^ embryos^18^. Together, these findings indicate that Runx1 and CBFβ play distinct but complementary roles in regulating hematopoiesis.

Runx1 interacts with various other transcriptional co-activators via its activation domain^19,20^, which leads to increased DNA binding and allows dynamic regulation of different genes involved in hematopoietic differentiation, ribosome biogenesis, cell cycle regulation, and p53 and transforming growth factor-β signaling pathways^21–24^. During EHT, Runx1 recruits the hematopoietic regulators SCL/TAL1 and FLI1 from ETS/GATA sites to RUNX1/ETS sites in hemogenic endothelial cells to activate the hematopoietic transcriptional program^25^. Runx1 binding to ETS transcription factors regulates a variety of genes via ETS activation on a Runx1/ETS composite DNA sequence^26^. Studies using genome-wide binding mapping in human CD34+ HSPCs revealed that a heptad of transcription factors (FLI1, ERG, GATA2, RUNX1, TAL1, LYL1, LMO2) showed distinct chromatin occupancy and enhancer-promoter interactions across cell types, which were associated with cell-type-specific gene expression^27,28^. Heptad-occupied regions in HSPCs were subsequently bound by lineage-defining transcription factors including PU.1 and GATA1 to direct cell fate decisions^27^. Furthering our understanding of the complex regulatory networks involved in HSC proliferation and differentiation may help to reveal new pathways for therapeutic targeting of HSC expansion and clonal diversity.

To identify new mechanisms regulating HSC clonality and expansion, we performed a chemical screen for modulators of Runx1 using zebrafish embryo cell cultures. We found that the Runx1/CBFβ inhibitor Ro5-3335 was a strong enhancer of *Runx1:eGFP* cells both in culture and *in vivo*. Ro5-3335, a benzodiazepine derivative, was first identified in a RUNX1–CBFβ interaction screen using both AlphaScreen and time-resolved fluorescence resonance energy transfer (TR-FRET) methods^29^. This chemical was shown to physically interact with both RUNX1 and CBFβ using a UV absorption depletion assay, with stronger binding to RUNX1 and weaker binding to CBFβ^29^. Notably, Ro5-3335 did not affect formation of the RUNX1–CBFβ complex in gel-shift or pull-down assays, indicating that it cannot fully disassociate the RUNX1–CBFβ complex but may induce a conformational change. Later, it was reported that Ro5-3335 had no effect on CBFβ-Runt domain binding and no interaction of the small molecule with either CBFβ or RUNX1 was detected by NMR^30^, indicating that it may have an alternative mechanism of action independent of RUNX1–CBFβ interaction. Ro5-3335 has also been reported to suppress tumor cell growth in leukemia^29^, pancreatic ductal adenocarcinoma^31^ and melanoma^32^, and shown to inhibit pathological neovascularization^33,34^, further suggesting that Ro5-3335 may modulate RUNX1 in multiple diverse ways.

Here, we discovered a new mechanism of action of Ro5-3335 in regulating HSC self-renewal and clonal diversity. To date, no chemical has been reported to increase the number of stem cell clones *in vivo*. In zebrafish, we showed that Ro5-3335 promotes HSPC divisions and increases HSC clonal diversity in adulthood. We also found increased clonal diversity in murine HSPC cultures treated with Ro5-3335. Using human CD34+ HSPCs and induced pluripotent stem cell (iPSC)-derived hemogenic endothelium, we showed that the effect of Ro5-3335 on HSPC expansion is conserved in human systems. Using both ChIP-seq and CUT&RUN analysis, we revealed that Ro5-3335 treatment drives a novel binding of ELF1/2 to the RUNX1 bound DNA regions, leading to increased transcription of cell cycle and hematopoietic target genes. *Elf2b* knockout in zebrafish abolished the effect of Ro5-3335 on HSPC expansion. Together, our studies reveal that the RUNX1 modulator Ro5-3335 alters RUNX1/ELF complex binding, leading to greater HSC production and a previously unknown mechanism that increases stem cell clonality.

## Results

### A zebrafish chemical genetic screen identifies Ro5-3335 as a compound that expands hematopoietic stem cells

We adapted our previously reported zebrafish embryo culture system^35,36^ to identify chemical modulators of HSPC expansion. Using the *Runx1:eGFP* transgenic reporter to specifically label HSPCs^37^, we dissociated zebrafish embryos into single cells at 24 hours post fertilization (hpf). Cells were plated with chemicals for 2 days until fluorescence was evaluated (Figure 1A). We screened 3,841 bioactive small molecules in duplicate and identified 22 chemical hits that induced *Runx1:eGFP* fluorescence (Table S1). Ro5-3335 scored as an inducer of *Runx1:eGFP* in our culture assay (Figure S1A). Previous studies show that Ro5-3335 affects definitive hematopoiesis, but the specific effect on Runx1-expressing HSCs themselves has not been evaluated^18^.

**Figure 1.**
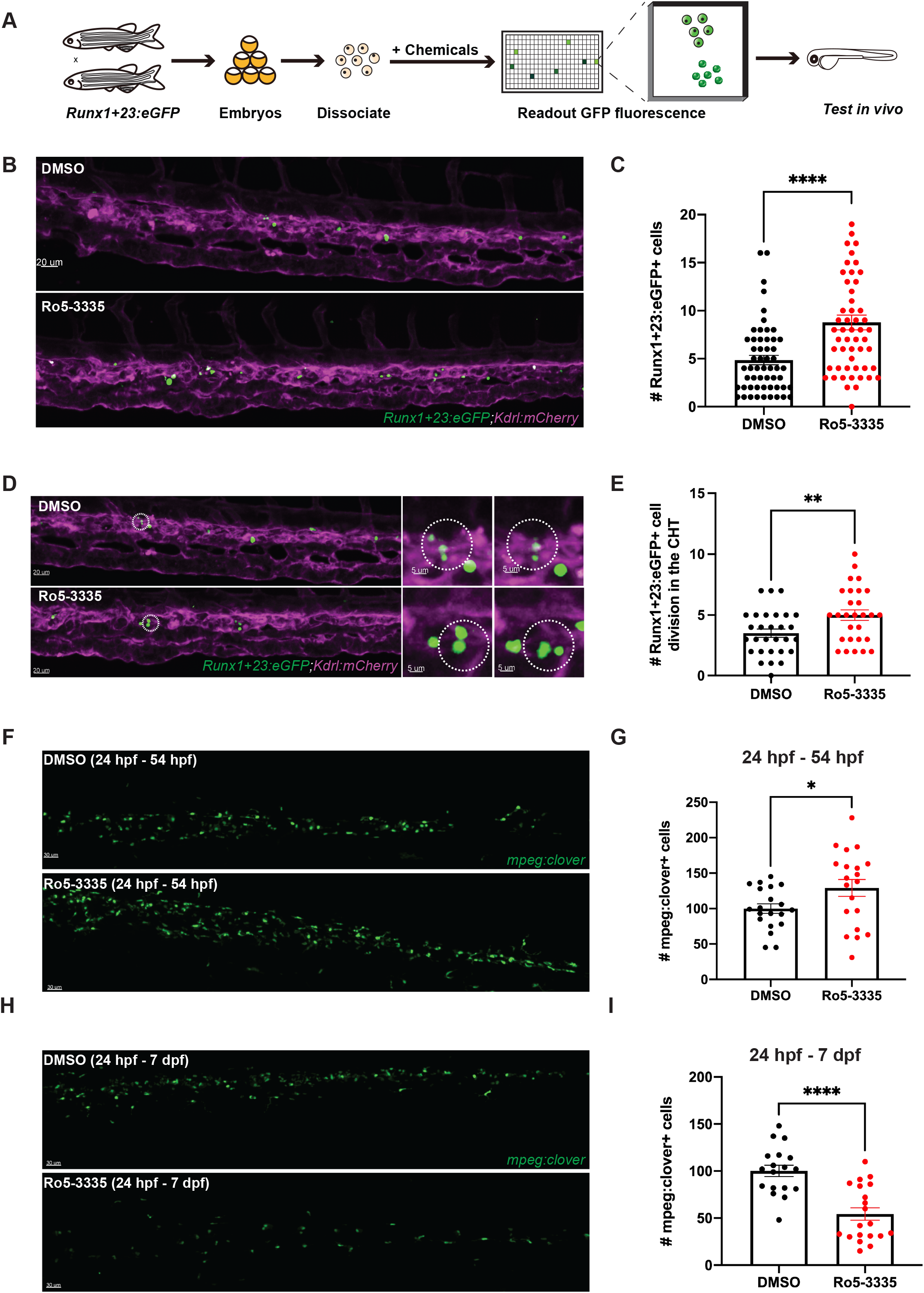
Ro5-3335 increases HSCs during definitive hematopoiesis. (A) Schematic of zebrafish embryo cell culture chemical screen. (B) Representative images of the CHT of *Runx1:eGFP;kdrl:mCherry* embryos treated with DMSO or Ro5-3335 from 24 hpf and imaged at 54 hpf. Scale bar, 20µm. (C) *Runx1:eGFP* embryos were treated with DMSO or Ro5-3335 from 24 hpf and *Runx1:eGFP+* in the CHT were quantified at 54 hpf (DMSO, n = 55; Ro5-3335, n = 54). (D, E) *Runx1:eGFP;kdrl:mCherry* embryos were treated with DMSO or Ro5-3335 from 24 hpf and time lapse imaging was performed from 45 to 57 hpf to quantify HSPC divisions. White dotted circles in (D) indicate a pair of dividing *Runx1:eGFP+* cells in the CHT. Scale bar, 20µm. Representative movies can be found in Movie S1-2 (DMSO, n = 29; Ro5-3335, n = 28). (F, G) *mpeg1:Clover* embryos were treated with DMSO or Ro5-3335 from 24 hpf to 54 hpf, and macrophages were quantified at 7 dpf (DMSO, n = 20; Ro5-3335, n = 20). Scale bar, 30µm. (H, I) *mpeg1:Clover* embryos were treated with DMSO or Ro5-3335 from 24 hpf to 7 dpf, and macrophages were quantified at 7 dpf (DMSO, n = 18; Ro5-3335, n = 20). Scale bar, 30µm. Bar graphs depict mean and SEM and each dot represents one fish. Statistical analyses were performed using two-tailed Student’s t-tests (*, P<0.05; **, P<0.01; ****, P <0.0001).

To further explore the effect of Ro5-3335 on HSCs, we quantified the number of *Runx1:eGFP*+ cells in live zebrafish embryos treated with Ro5-3335 between 24 and 54 hpf. During this time window, the majority of stem cells are born in the AGM and migrate via circulation to the intermediate niche in the caudal hematopoietic tissue (CHT) where they divide, before ultimately making their way to the kidney marrow from around 4.5 days post fertilization (dpf). After Ro5-3335 treatment, we found a significant increase in HSCs in the zebrafish CHT at 54 hpf, in line with the effect observed in our culture assay (Figure 1B, C). We deduced that this expansion could be explained either by enhanced specification of nascent HSCs in the AGM resulting in the arrival of more HSCs in the niche, or by more frequent stem cell divisions in the CHT. To examine HSC specification, we performed high resolution live imaging of HSPC emergence from the AGM in *cmyb:eGFP;kdrl:mCherry* embryos between 30 and 48 hpf, which showed that the number of budding HSCs was unaltered in the presence of Ro5-3335 (Figure S1B, C). Live imaging of the CHT in *Runx1:eGFP;kdrl:mCherry* embryos revealed instead that HSCs undergo a significantly increased number of divisions between 45 and 57 hpf upon Ro5-3335 treatment (Figure 1D, E, Movie S1-2), which we estimate would most likely correlate to one additional cell division. Taken together, these data suggest that Ro5-3335 regulates HSC expansion, without affecting early HSC emergence.

We next hypothesized that the increased HSCs might give rise to expanded differentiated cell types. Using a transgenic zebrafish line expressing the fluorescent protein NLSmClover driven by the *mpeg1* promoter (*mpeg1:Clover*) to label macrophages^38^, we investigated the effect of Ro5-3335 treatment from 24 hpf to 7 dpf to capture macrophages from the definitive wave of hematopoiesis^39^. When embryos were treated with Ro5-3335 from 24 to 54 hpf, the number of macrophages at 7 dpf was significantly higher than the controls (Figure 1F, G). In contrast, when embryos were continuously treated with Ro5-3335 from 24 hpf up to 7 dpf, we found that the number of macrophages present in the whole embryo at 7 dpf was dramatically reduced in the presence of Ro5-3335 compared to controls (Figure 1H, I). This indicated that prolonged exposure to Ro5-3335 might not only induce the division of HSCs but also block their differentiation. With only transient exposure, the cells can resume differentiation once Ro5-3335 is removed. The resulting increase in macrophages we observed suggests that blocking differentiation for a brief period allows HSCs to undergo addition self-renewing divisions, creating a larger pool of stem cells that may consequently give rise to expanded populations of downstream progeny.

### Ro5-3335 increases HSC clone number and improves HSC function upon transplantation

Our previous studies have demonstrated that environmental stimuli and cellular interactions occurring within the CHT niche during embryonic development have a profound effect on establishing the number of HSC clones that contribute to adult hematopoiesis^6,7^. To date, no chemical has been reported to increase the number of stem cell clones *in vivo*. Given that Ro5-3335 caused an increase in HSPC expansion in the CHT, we sought to investigate if this would translate to changes in HSC clonality in the adult kidney marrow. We used the Zebrabow HSC color barcoding system^6,40^ to uniquely label each individual HSC clone by exposing embryos to 4-hydroxytamoxifen (4-OHT) at 24 hpf, in combination with Ro5-3335 or DMSO treatment until 54 hpf (Figure 2A). The drl:creER^T2^ driver is active in early hematopoiesis^6^, and color labeling at 24 hpf, immediately prior to HSC emergence, ensured that all new clones emerging in the AGM would receive a unique and heritable color barcode. Thorough washing of embryos at 54 hpf guaranteed that HSPCs were only transiently exposed to Ro5-3335, so that long-term differentiation was not affected. To evaluate HSC clonality, we raised the embryos for 3 months then analyzed the kidney marrow (the adult hematopoietic niche, equivalent to the mammalian bone marrow) by flow cytometry. We then performed clonal analysis on cells from the myeloid compartment, as previously described^6,7^. We found that transient exposure to Ro5-3335 during a short time window of development was sufficient to significantly increase HSC clone number in the adult marrow (Figure 2B-D). Using the Gini coefficient of each clone as a measure of clonal dominance^10^, we confirmed that there were no dominant clones in the Ro5-3335 treated cohort (Figure S2A), but rather Ro5-3335 induced development of more clones that were smaller in size compared to the DMSO cohort (Figure S2B). To our knowledge, this is the first time that pharmacological manipulation has resulted in an increase of HSC clones in an *in vivo* setting.

**Figure 2.**
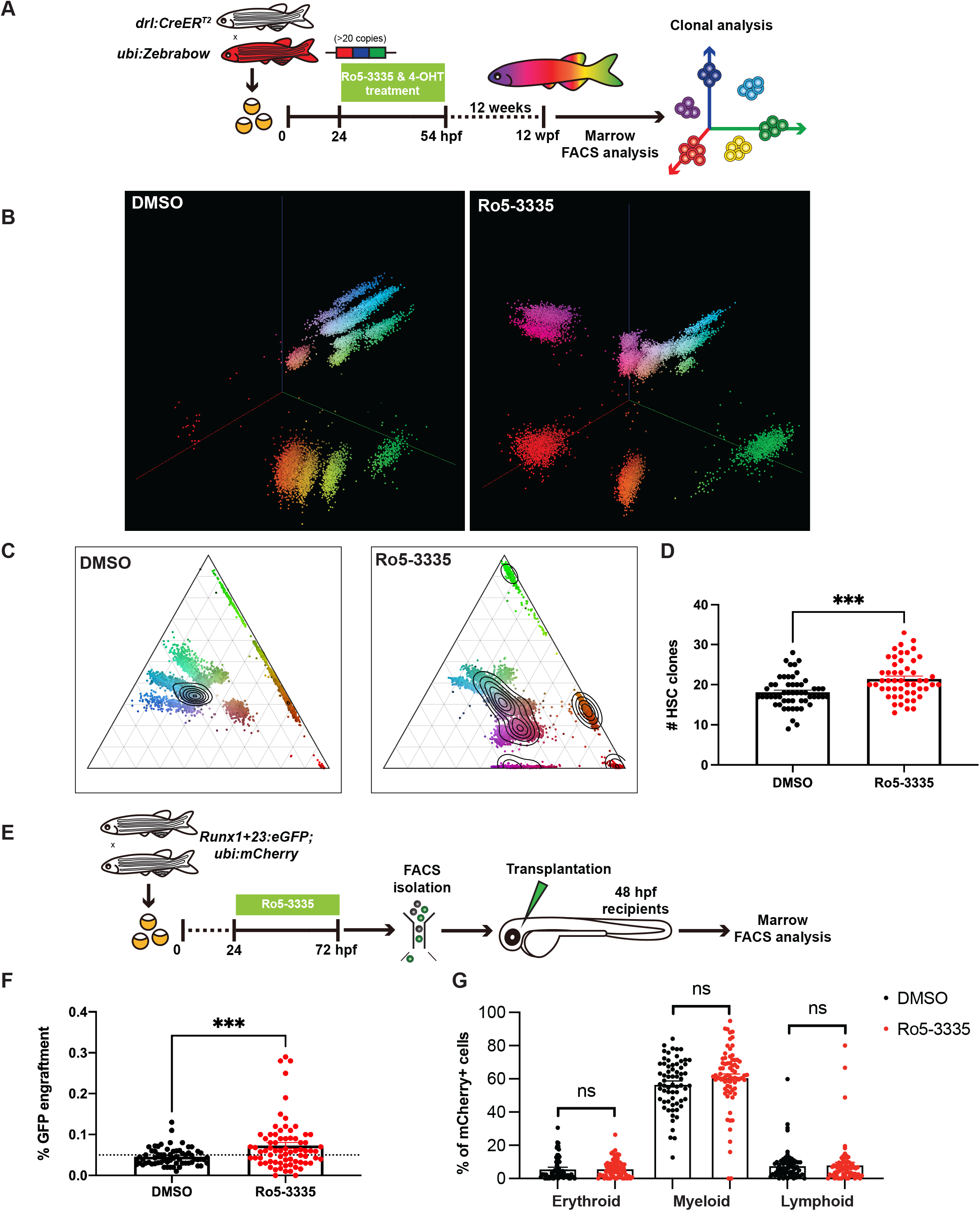
Ro5-3335 increases HSC clone number and improves HSC engraftment. (A) Schematic of Zebrabow HSC color labeling system. (B-D) *ubi:Zebrabow* and *drl:CreER^T2^* embryos were treated with DMSO or Ro5-3335 along with 4-OHT from 24 to 54 hpf then raised to adulthood. Kidney marrow was analyzed at 12 wpf by FACS to quantify HSC clone number and clone size. (B) Representative 3D cluster plots and (C) corresponding ternary plots showing color distribution in granulocytes. (D) Quantification of HSC clone number (DMSO, n = 54; Ro5-3335, n = 52). (E) Schematic of embryo-to-embryo transplantation assay. (F) Engraftment of *Runx1:eGFP* cells shown as % of total live cells from the kidney marrow at 3-4 months post-transplant. Dotted line indicates threshold for positive engraftment above background (0.05%) (DMSO, n = 65; Ro5-3335, n = 73). (G) Assessment of contribution to erythroid, myeloid and lymphoid lineages based on *ubi:mCherry*+ cells shown as % of total live cells (DMSO, n = 65; Ro5-3335, n = 73). Bar graphs depict mean and SEM and each dot represents one fish. Statistical analyses were performed using two-tailed Student’s t-tests (***, P<0.001; ns = non significant).

To further characterize the expanded population of HSPCs that arise with Ro5-3335 stimulation and investigate whether they still have the potential to function as *bona fide* stem cells, we performed embryo-to-embryo transplantation experiments (Figure 2E). Based on our previous approach^37^, we isolated double transgenic *Runx1:eGFP+;ubi:mCherry*+ HSPCs from Ro5-3335 treated embryos and transplanted equal numbers of *Runx1:eGFP+;ubi:mCherry*+ HSPCs into *casper* embryo recipients at 48 hpf. The thymus is not fully formed at this stage, allowing for introduction of exogenous cells without immune rejection, and presence of the *ubi:mCherry* transgene allowed us to label the progeny of the donor *Runx1:eGFP* cells. At ∼3 months post-transplant, engraftment was assessed by FACS analysis of the kidney marrow, which revealed that *Runx1:eGFP+* HSPCs from Ro5-3335 treated embryos could engraft more effectively than those isolated from vehicle control embryos (Figure 2F). In addition, we observed that these engrafted cells had the capacity to undergo multilineage differentiation, evidenced by the presence of *ubi:mCherry+* cells within each lineage compartment without lineage skewing (Figure 2G). Next, we performed secondary transplantation experiments by isolating *Runx1:GFP*+ cells from the kidney marrow of our primary recipients at 3 months post-transplant and retransplanting equal numbers of *Runx1:eGFP+;ubi:mCherry*+ HSPCs into *casper* embryo recipients at 48 hpf. At 3 months post-secondary transplant, GFP+ cells originating from both the DMSO and Ro5-3335 treated cells could be observed in the marrow (Figure S2C), indicating that they retain their long-term self-renewal activity. Taken together, these data indicate that Ro5-3335 expands HSPCs *in vivo* by increasing the number of clones formed during early development and that these cells maintain their stem cell characteristics through adulthood in terms of engraftment, self-renewal and differentiation.

### Ro5-3335 increases clonality and self-renewal in mouse HSCs ex vivo

We next sought to validate the role of Ro5-3335 in the regulation of clonality and self-renewal in mouse HSCs. We barcoded individual HSCs using the LARRY (Lineage and RNA recovery) barcoding system^41^ and tracked the clonal expansion of each HSC clone in the presence of DMSO or Ro5-3335 for 14 days (Figure 3A). We integrated DMSO and Ro5-3335 samples with day 10 and day 14 WT polyvinyl alcohol (PVA)-based HSC cultures^41^ for unbiased cell clustering (Figure 3B, S3A, B), and successfully detected HSCs, multipotent progenitors (MPPs), granulocyte-monocyte progenitor (GMPs) and megakaryocytes (Mk) in the culture in both the DMSO and Ro5-3335 conditions (Figure 3C). We recovered an increased number of HSC clones in the Ro5-3335 treated HSC cultures (Figure 3D), suggesting that Ro5-3335 enhanced HSC self-renewal in *ex vivo* culture. We further assessed HSC clonal output by analyzing the cellular distribution patterns between HSC and non-HSC clusters as previously described^41^. Ro5-3335 treated HSCs showed a decreased output activity, suggesting that Ro5-3335 stimulates self-renewal while inhibiting differentiation (Figure 3E). Both cluster-based (Figure 3F) and Milo-based (Figure 3G) differential abundance analysis revealed a significant reduction in committed lineages (Flt3+GMPs, GMPs, and Mks-) and an enrichment of Flt3+HSCs in Ro5-3335 samples compared to DMSO controls. Single cell transcriptomics indicated that Ro5-3335 treated HSCs upregulated pathways related to HSC proliferation and increased metabolic activity, such as oxidative phosphorylation. Meanwhile, pathways related to inflammation (TNFα and IFNγ) were downregulated (Figure S3C-E). Overall, these results support that Ro5-3335 increases murine HSC self-renewal and clonality while inhibiting differentiation, in line with our findings in zebrafish.

**Figure 3.**
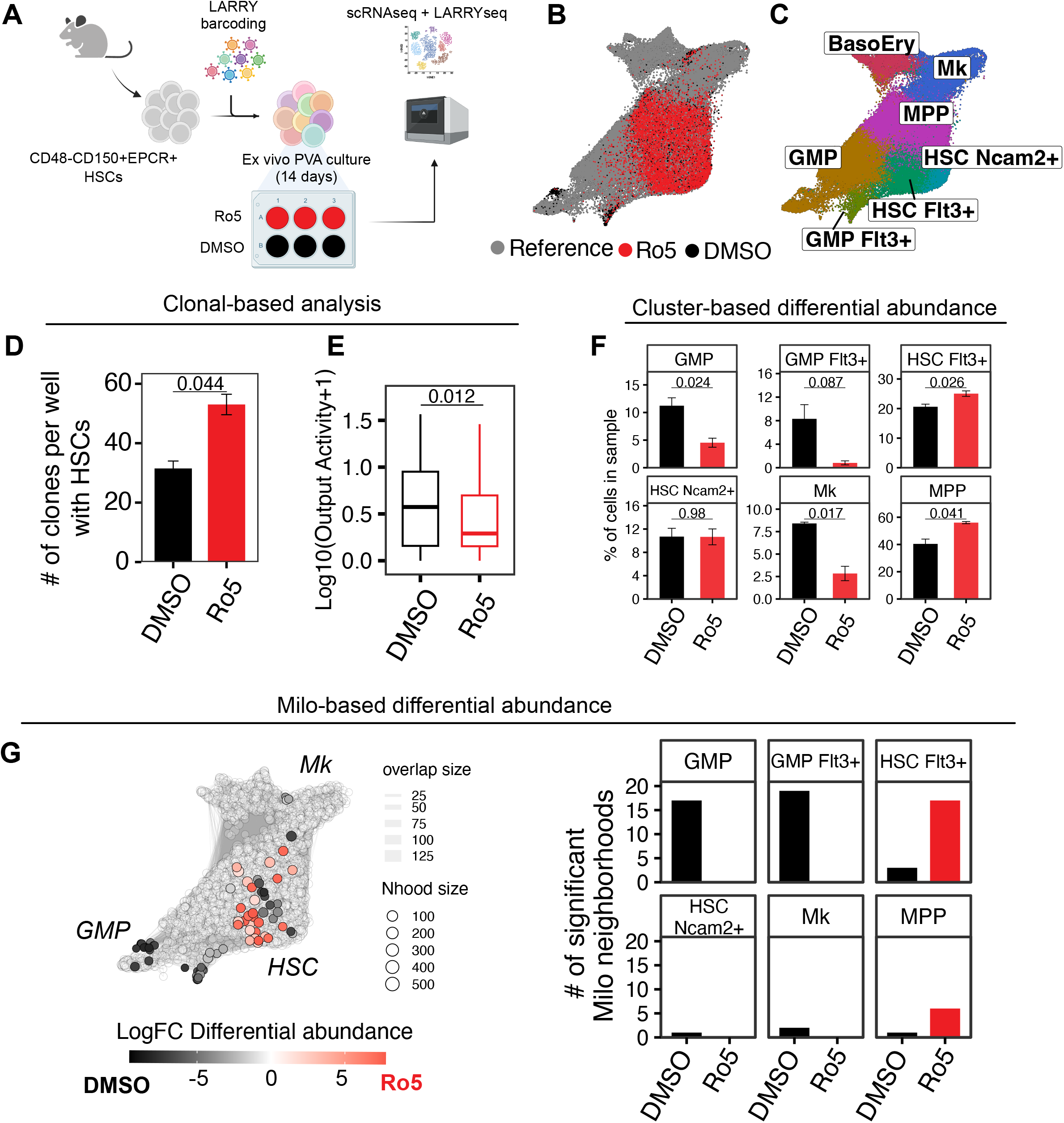
Ro5-3335 increases clonality and self-renewal in murine HSC cultures. (A) Schematic of experimental design. Cultures were started using 10,000 E-SLAM HSCs (lineage−Sca-+cKit+CD48−CD150+CD201+), transduced with the LARRY barcoding system -MOI 0.3-expressing the T-Sapphire fluorescent protein. Labeled HSCs were isolated by FACS then split equally into DMSO and Ro5-3335 containing wells (n = 3). After 14 days of culture, cells were profiled by scRNA-seq and LARRY-seq. (B) UMAP of DMSO control (blue), Ro5-3335 (yellow) cells and reference cells (grey). (C) Duplicate of UMAP in (B) with cells colored based on their associated cell type. (D) Number of clones per well detected in DMSO and Ro5-3335 samples. Data is represented as mean +/− SD. Wilcox-rank sum test, n = 2. (E) Boxplots representing the distribution of Output Activity (self-renewal vs differentiation) of DMSO and Ro5-3335 clones. Wilcox-rank sum test, n = 118 (DMSO) and 313 (Ro5-3335). (F) Cluster-based differential abundance analysis. Bars represent the percentage of cells associated to control DMSO or Ro5-3335 treated cells. Bar graphs depict mean and SD. Wilcox-rank sum test, *n* = 2. (G) Left: Milo-based differential abundance. UMAP representing Milo neighborhoods colored based on the Log2FC differential abundance. Blue colored neighborhoods are enriched in DMSO samples, yellow indicates enrichment in Ro5-3335. Grey neighborhoods are not significantly enriched in any condition (FDR < 0.1). Right: Number of significant Milo neighborhoods per cell type in DMSO and Ro5-3335. These are the same neighborhoods represented in the UMAP grouped by cell type.

### Ro5-3335 enhances human HSC expansion ex vivo

We next assessed Ro5-3335 function in human *in vitro* systems. To examine whether Ro5-3335 increases HSC production from induced pluripotent stem cells (iPSCs), human iPSCs were induced into hemogenic endothelial cells by embryoid body formation and then differentiated to hematopoietic progenitors through EHT (Figure 4A). Hemogenic endothelial cells were treated with Ro5-3335 on day 0 of EHT. At the middle stage of EHT (day 4), both adherent and floating cells were harvested and progress of EHT was analyzed by flow cytometry for CD34 and CD45 expression. Two independent dose-test experiments showed that Ro5-3335 treatment increased the percentage of CD34+ cells while decreasing acquisition of CD45 positivity, resulting in a reduction of CD34+CD45+ double positive hematopoietic progenitors (Figure 4B, C). We observed fewer cells in the iPSC culture with high doses of Ro5-3335 (25 µM), suggesting toxicity at a high dose despite high efficacy. By day 7 of EHT, cells in control conditions reach the CD45+ single positive state, at which stage they have lost their ability to contribute to multiple hematopoietic lineages. Ro5-3335 treatment maintained hematopoietic cells in a less committed state, increasing the proportion of multipotent hematopoietic cells (Figure S4A-D), and suggesting that Ro5-3335 delayed CD45 acquisition but did not stop it. These data support the observation we made in our zebrafish and murine assays, in which we detected a block in differentiation that may allow increased time for cell cycling and HSC expansion.

**Figure 4.**
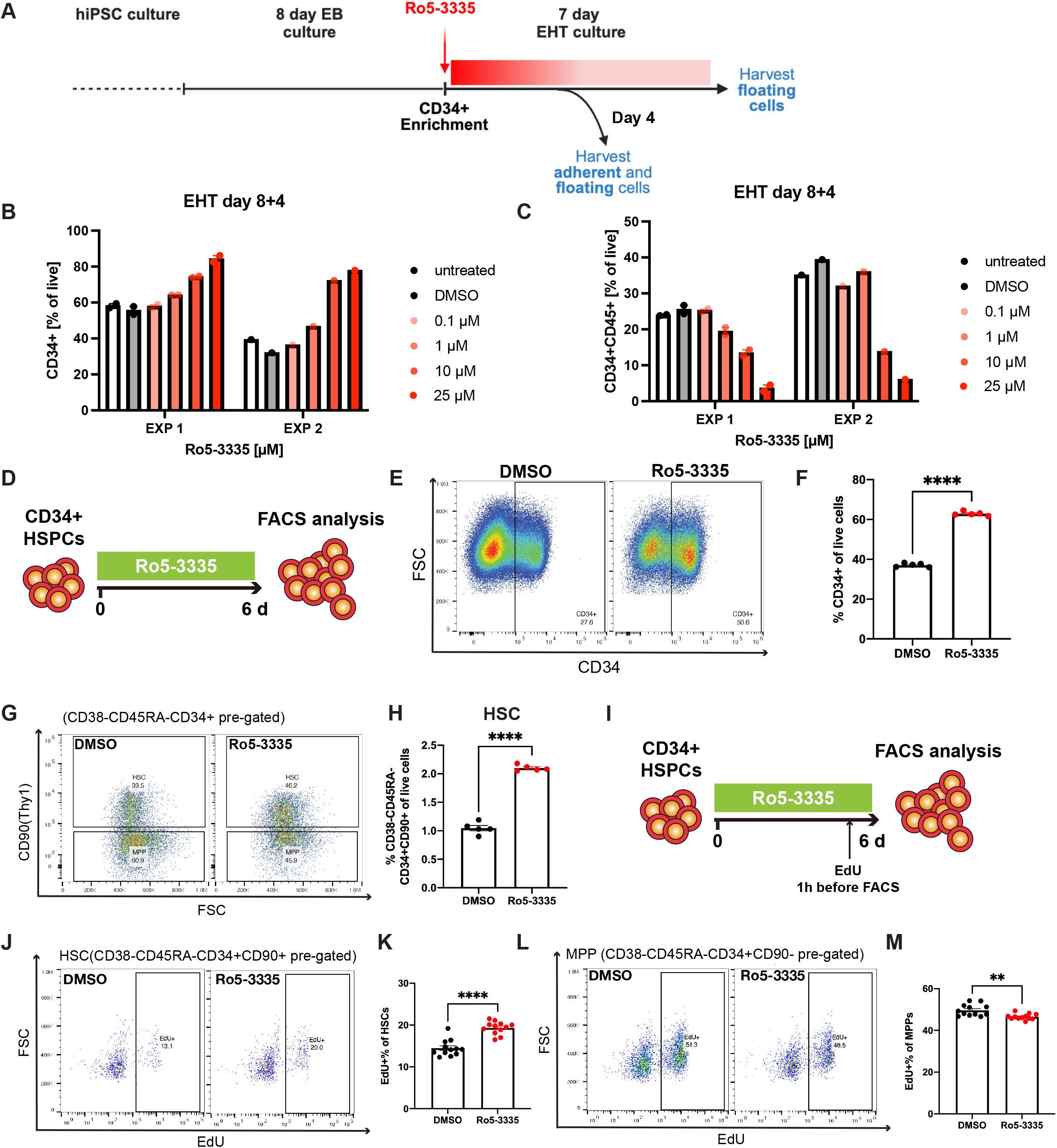
Ro5-3335 enhances human HSC expansion *ex vivo*. (A) Schematic of hiPSC-derived HSPC culture. Hemogenic endothelial cells were MACS enriched for CD34+ on day 8 of embryoid body (EB) culture and plated in EHT conditions in the presence of Ro5-3335 or DMSO. EXP1 and 2 indicate two independent experiments. Flow cytometric quantification of (B) CD34+ single positive and (C) CD34+CD45+ double positive cells in adherent and floating cells at day 4 of EHT-culture (n = 2). (D-H) Human CD34+ HSPCs from mobilized peripheral blood were expanded for 6 days and cells were harvested for FACS analysis at day 6. Cells were continuously treated with DMSO or Ro5-3335 in culture. (D) Schematic of CD34+ HSPC culture. (E) FACS plot and (F) quantification of CD34+ cells at day 6. (G) FACS plot and (H) quantification of HSCs (CD38-CD45RA-CD34+CD90+) at day 6 (n = 5). (I-M) EdU was added to culture media 1 hour before harvest for FACS and EdU analysis at day 6. (I) Schematic of the CD34+ HSPC culture for EdU analysis. (J) FACS plot and (K) quantification of EdU incorporation of HSCs (CD38-CD45RA-CD34+CD90+) at day 6. (L) FACS plot and (M) quantification of EdU incorporation of MPPs (CD38-CD45RA-CD34+CD90-) cells at day 6 (DMSO, n = 5; Ro5-3335, n = 5). Bar graphs depict mean and SEM. Statistical analyses were performed using two-tailed Student’s t-tests (**, P<0.01; ****, P <0.0001).

To further investigate whether Ro5-3335 increases HSC expansion *ex vivo*, human mobilized peripheral blood (mPB) CD34+ HSPCs were cultured in expansion media with Ro5-3335. At day 6 of expansion, the cell culture was analyzed by flow cytometry to examine the proportion of CD34+ progenitors, HSCs (CD38-CD45RA-CD34+CD90+), and MPPs (CD38-CD45RA-CD34+CD90-) (Figure 4D). We observed an increased percentage of CD34 expressing cells in cells from day 6 of the expanded HSPC culture (Figure 4E, F), suggesting that Ro5-3335 maintains the progenitor state and prevents differentiation, in line with our previous data. Total CD34+ progenitor cells did not change significantly (Figure S4E), which might be due to the slight decrease in total cell numbers, indicating that Ro3-3335 limits the expansion of progenitor cells. Ro5-3335 increased the percentage of HSCs (Figure 4G, H) but decreased the percentage of MPPs (Figure S4F), suggesting that Ro5-3335 specifically promoted the expansion of HSCs but not MPPs. Furthermore, we observed that Ro5-3335 treatment promoted HSC expansion from 15.8 fold to 20.3 fold (Figure S4G), while inhibiting MPP expansion from 35.6 fold to 15.6 fold (Figure S4H). To determine whether Ro5-3335 expands HSCs by promoting proliferation as we had observed *in vivo*, we added thymidine analog ethynyl deoxyuridine (EdU) to the media to label dividing cells 1 hour before harvest (Figure 4I). We examined the EdU incorporation rate by flow cytometry and observed that more HSCs were EdU+ (Figure 4J, K), indicating that Ro5-3335 promoted HSC proliferation in *ex vivo* culture. In contrast, MPPs were generally more proliferative than HSCs under control conditions, but Ro5-3335 decreased their proliferation (Figure 4L, M). These findings revealed that Ro5-3335 specifically promotes HSC but not MPP division, providing additional evidence for a block in differentiation. Altogether, our studies showed that Ro5-3335 enhanced HSC expansion *ex vivo* and suggest an important role for Ro5-3335 in maintaining HSC stemness.

### ELF factors govern the RUNX1-mediated transcriptional response upon Ro5-3335 treatment

The transcription factor RUNX1 is an essential regulator of hematopoiesis and controls the activation or repression of specific genes by binding to DNA along with an array of cofactors that modulate its transcriptional activity^42^. One of these cofactors, CBFβ, is known to relieve RUNX1 auto-inhibition and stabilize the interaction with its DNA binding motif^13,15,16^. Pharmacological disruption of the interaction between RUNX1 and CBFβ, for example as reported with Ro5-3335^29^, would predict reduced RUNX1 occupancy and altered expression of target genes. However, the mechanism of Ro5-3335 may be more complex, and other studies have indicated that Ro5-3335 does not directly inhibit RUNX1/CBFβ binding but rather functions via an indirect mechanism involving an alternative cofactor^30^. Indeed, when we tested two inhibitors that bind directly to CBFβ to inhibit interaction with RUNX1, we did not detect an increase in *Runx1:eGFP* HSCs in our zebrafish assay (Figure S5A). This suggests that the response we observe on HSC expansion in the presence of Ro5-3335 is likely not due to disruption of Runx1/Cbfβ binding, but rather another mechanism likely involving additional transcriptional regulators is at play.

To understand the mechanism for increased HSC expansion by Ro5-3335, we performed ChIP-seq and CUT&RUN on human CD34+ HSPCs. We investigated whether Ro5-3335 could induce changes in DNA occupancy of RUNX1-bound genes by ChIP-seq profiling of RUNX1 and CBFβ in CD34+ HSPCs following treatment with Ro5-3335 or DMSO for 24 hours, to detect the immediate DNA binding changes. Under the same conditions, we additionally performed ChIP-seq for H3K27-acetylation marks (H3K27ac) and ATAC-seq to investigate the Ro5-3335-mediated change in genome-wide activity and chromatin accessibility of enhancers. Strikingly, we observed limited changes to the enhancer landscape and DNA binding of CBFβ, but the DNA occupancy of RUNX1 increased considerably upon Ro5-3335 treatment (Figure 5A). This suggested that additional cofactors may be involved to mediate RUNX1 DNA binding activity. Chromatin accessibility and H3K27-acetylation also showed limited changes, indicating that these RUNX1 bound sites are already open and active.

**Figure 5.**
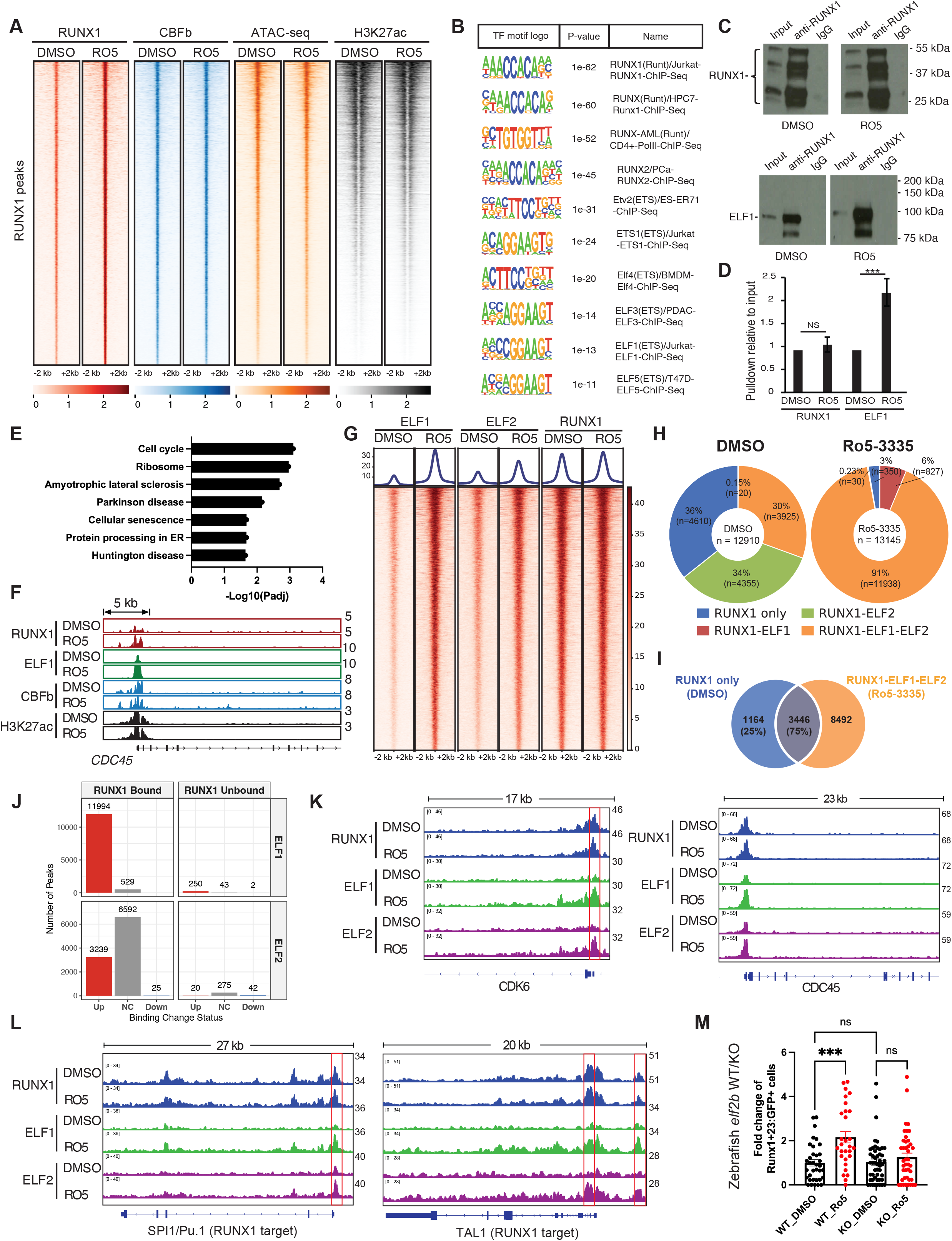
ELF factors govern the RUNX1-mediated response upon Ro5-3335 treatment. (A) Peak density plots of RUNX1 ChIP-seq, CBFβ ChIP-seq, ATAC-seq, and H3K27ac ChIP-seq of CD34+ HSPC cultures treated with DMSO or Ro5-3335 for 24 h. (B) Motif analysis of genes with increased RUNX1 binding upon Ro5-3335 treatment. Genes were pre-selected to have RUNX1-CBFβ co-binding. (C-E) CD34+ HPSCs were expanded for 6 days and then treated with DMSO or Ro5-3335 for 24 h. Cells were harvested for RUNX1 co-immunoprecipitation (co-IP). (C) Western blot plot of RUNX1 protein and ELF1 protein from RUNX1 co-IP. (D) Quantitative analysis of the pulldown relative to input of RUNX1 protein and ELF1 protein from RUNX1 co-IP. (E) KEGG analysis of RUNX1+ELF1 co-bound genes showed increased RNA expression and increased RUNX1 binding with Ro5-3335 treatment versus DMSO. (F) Representative track view of peaks from RUNX1 ChIP-seq, ELF1 ChIP-seq, CBFβ ChIP-seq, and H3K27ac ChIP-seq of cell cycle gene CDC45. (G) Peak density plots of ELF1, ELF2, and RUNX1 CUT&RUN sequencing of CD34+ HSPC cultures treated with DMSO or Ro5-3335 for 24 h. (H) Donut chart analysis of RUNX1 sites showing RUNX1 co-occupancy with ELF1 and ELF2 with Ro5-3335 treatment versus DMSO. (I) Venn diagram analysis showing the number of RUNX1 sites that were RUNX1-bound only in DMSO and RUNX1/ELF1/ELF2 co-bound in Ro5-3335 conditions. The overlapping region in the Venn diagram indicates the regions that switched from RUNX1-bound only to RUNX1/ELF1/ELF2 co-bound upon Ro5-3335 treatment. (J) Number of ELF1/2 bound peaks showing differential binding (|Log2Fold Change| > 0.68) upon Ro5-3335 treatment. Peaks were further grouped by RUNX1 binding (NC, no change). (K-L) Representative track view of peaks from RUNX1, ELF1, and ELF2 CUT&RUN-seq analysis of representative (K) cell cycle genes CDK6, CDC45 and (L) known RUNX1 targets in hematopoiesis SPI1/Pu.1 and TAL1. (M) Embryos from *Elf2b+/−; Runx1:eGFP* incrosses were collected and treated with DMSO or Ro5-3335 from 24 hpf. *Runx1:eGFP+* cells in the CHT were quantified at 54 hpf. Each embryo was genotyped after imaging. (WT DMSO-treated, n = 35; WT Ro5-3335-treated, n = 29; KO DMSO-treated, n= 48; KO Ro5-3335-treated, n = 42). Bar graphs depict mean and SEM and each dot represents one fish. Statistical analyses were performed using one-way ANOWA test (***, P <0.001; ns = non significant).

To determine additional cofactors in the RUNX1/CBFβ complex that may lead to the increased occupancy of RUNX1 upon Ro5-3335 treatment, we performed transcription factor motif analysis on the RUNX1 and CBFβ co-bound genes that showed increased RUNX1 binding and increased RNA expression (log_2_ fold change, Ro5-3335 vs DMSO ≥ 0.5). We discovered an increased enrichment of ETS/ELF transcription factor motifs (Figure 5B), suggesting that ELF transcription factors may promote enhanced binding of RUNX1 at these enhancers to increase the expression of associated genes. We performed RUNX1 co-immunoprecipitation (co-IP) and confirmed the increased interaction between RUNX1 and ELF1 in CD34+ HSPCs upon Ro5-3335 treatment (Figures 5C, D). We then compared the expression of genes that showed increased co-binding of RUNX1 and ELF1 to the expression of genes that showed an increase in RUNX1 binding only with Ro5-3335 stimulation (Figure S5B). This analysis indicated that Ro5-3335 leads to moderate but significantly increased expression of RUNX1 and ELF1 co-bound genes compared to genes that show an increase in RUNX1 binding only. We next performed gene ontology analysis on the RUNX1 and ELF1 co-bound genes with increased RNA expression upon Ro5-3335 treatment and found that the cell cycle was the top altered gene set (Figure 5E), including genes such as CDC45 (Figure 5F) and CDC37L1 (Figure S5C). We further performed CUT&RUN analysis on ELF family members (ELF1 and ELF2) and RUNX1 on HSPCs and observed a global increase in ELF1/2 binding to RUNX1 sites with Ro5-3335 treatment (Figure 5G). Upon further examination of co-occupancy, we identified that RUNX1/ELF1/ELF2 co-bound sites increased from 30% (n=3,925) to 91% (n=11,938) in the presence of Ro5-3335 (Figure 5H). Strikingly, 75% (n=3,446) of the RUNX1-only sites in the DMSO control became co-bound with RUNX1/ELF1/ELF2 upon Ro5-3335 treatment (Figure 5I), indicating a novel recruitment of ELF1/2 to the RUNX1 complex, induced by Ro5-3335. ELF1 and ELF2 showed a global increase in DNA binding and most of the regions with increased ELF1/2 binding are indeed RUNX1-bound sites (Figure 5J), indicating a preferential recruitment of ELF1/2 to the RUNX1 complex. We observed that a number of cell cycle genes (CDK6, CDC45, CCND2, CDC37L1), along with RUNX1 targets that are key hematopoiesis regulators (SPI1/Pu.1, TAL1), showed increased ELF1/2 binding with Ro5-3335 treatment (Figure 5K, L and Figure S5E). Altogether, our data suggest that the increased recruitment of ELF factors to the RUNX1/CBFβ complex mediated by Ro5-3335 might induce transcriptional activation of cell cycle genes, which may serve as one of the mechanisms to promote HSPC expansion.

To further validate the ELF-dependent mechanism of Ro5-3335, we returned to the zebrafish model, which expresses 4 ELF orthologs; *elf1*, *elf2a*, *elf2b* and *elf3*^43^. In zebrafish embryos, the effect of Ro5-3335 on HSC expansion in the CHT was lost upon knockdown of *elf2b*, with HSC numbers returning to a level comparable to the DMSO control (Figure S5F). We generated an *elf2b* stable knockout (KO) mutant zebrafish using CRISPR/Cas9 technology and confirmed that the effect of Ro5-3335 on HSC expansion in the CHT was abolished with *elf2b* KO compared to the wild-type sibling control (Figure 5M, Figure S5G). The ETS domain of zebrafish Elf2b protein shares almost 100% homology with the ETS domain of human ELF family members (Figure S5H) and is the key domain for DNA binding. This further supports that the mechanism of Ro5-3335 may rely on increased interaction of Runx1 and Elf family members, and the subsequent transcriptional activation of a unique set of downstream target genes, to result in enhanced HSC expansion.

## Discussion

Hematopoiesis is tightly regulated by a balance between self-renewal, quiescence, and differentiation of HSCs. A defined number of HSC clones are born during early embryonic development, and these cells are responsible for sustaining the entire blood system throughout life. Clonal dynamics and cell fate decisions are controlled by extrinsic factors such as hypoxia, cytokines and ROS, and intrinsic factors including transcription factors, cell cycle regulators and epigenetic modifiers, both in normal hematopoiesis and hematopoietic malignancies^44^. The transcription factor RUNX1 is essential for definitive hematopoiesis and binds to DNA with cofactors including CBFβ to activate expression of hematopoietic target genes^38^. Genetic mutations in *runx1* cause defects in definitive hematopoiesis and fewer HSC clones remain to contribute to adult blood^9,10^. Though this and other mechanisms that reduce the diversity of HSC clones have been explored^6,7^, little is known about the mechanisms that promote clonal diversity.

Here, we identified a new way to chemically modulate clonality during embryonic development to result in an expanded number of HSC clones in adulthood. These HSCs undergo more frequent divisions in the zebrafish CHT and expansion in *ex vivo* mouse HSC cultures. These findings were conserved in human CD34+ cells, where we observed an increase in HSCs that were EdU+. Our combined approaches across multiple model systems suggest that Ro5-3335 retains HSCs in a less differentiated state, allowing additional time for self-renewal divisions, and blocking further differentiation. Cell cycle studies using the label-retaining reporter H2B-GFP to trace symmetric self-renewal divisions over time indicate that only 1% of repopulating cells within the aging HSC compartment are long term HSCs^45^. Ro5-3335 may extend these self-renewal divisions to delay the loss of HSC stemness during ageing. This is supported by our observations that Ro5-3335 blocks differentiation of iPSC-derived hematopoietic progenitors, CD34+ progenitors, murine HSCs and zebrafish macrophages. Other reports have also shown that Ro5-3335 can block myeloid-skewed hematopoiesis^46^. Furthermore, we found that HSPCs isolated from zebrafish embryos treated with Ro5-3335 showed higher chimerism upon transplantation, and this engraftment potential was maintained upon secondary transplantation. Our data suggest that Ro5-3335 enhances the fundamental properties of HSPCs to direct them towards a more ‘stem-like’ fate. Interestingly, Ro5-3335 showed a minor increase in MPPs (p = 0.041) in the mouse PVA-culture system but significantly decreased MPP expansion in the human CD34+ HSPC culture system. This selective effect might be due to the different differentiation state of MPPs between the two culture systems, with MPPs in murine cultures perhaps being in a more primitive state. This indicates a potential need for improvement of the standard human CD34+ HSPC expansion culture system to better maintain stemness, and Ro5-3335 may be a good candidate for this.

Combinatorial transcription factor interactions allow precise control over cell fate. A heptad of transcription factors (SCL/TAL1, LYL1, LMO2, GATA2, RUNX1, ERG and FLI-1) elegantly regulate normal HSC development^28^. Integrating the cell-type specific transcription factors MITF and c-FOS with the heptad factors allows mast cell specific gene expression^47^, while replacing GATA2 with GATA1 permits expression of erythroid related genes^48^. The addition or replacement of lineage determining transcription factors remodels the transcription complex, resulting in genome-wide alterations in binding sites and binding affinity. This allows for complex regulation and fine-tuning of gene expression involved in cellular activities and cell fate determination. Our work suggests that Ro5-3335 remodels the RUNX1 transcription complex to allow ELF1/2 binding and specific expression of cell cycle related target genes that enhance HSC self-renewal and prevent differentiation. The precise molecular mechanism of Ro5-3335 is unclear, though it has been reported to disrupt the RUNX1-CBFβ binding interaction by inducing a conformational change in the protein complex that may increase the distance between RUNX1 and CBFβ^29^. It is possible that this structural change might expose novel binding sites to allow additional factors, including ELF/ETS family members, to bind the complex and activate transcription of target genes for HSC self-renewal. When we tested other RUNX1-CBFβ inhibitors that were validated by FRET, NMR and co-IP^30^, we did not see the same effect on HSC expansion *in vivo*. This provides further evidence that Ro5-3335 is acting via a mechanism independent of CBFβ. It has also been suggested to inhibit SMARCA2, a member of the SWI/SNF chromatin remodeling complex, which is reported to interact with RUNX1 to control hematopoietic specific gene expression^49^. Further investigation is required to define the precise molecular target of Ro5-3335 and how this regulates the transcriptional program to activate target genes that control HSC self-renewal.

Though we identified ELF1/2 binding to RUNX1 as a potential mechanism to promote HSC stemness in human CD34+ cells following Ro5-3335 treatment, we did not detect a loss of HSPC expansion when we knocked out *elf1* in zebrafish. The zebrafish Elf1 protein shares a similar level of homology (43.3%) to human ELF1 protein as Elf2b; the knockout of which was sufficient to block the effect of Ro5-3335. We searched the ZFIN expression database^50,51^ and found that *elf1* is not detected in zebrafish embryos at 48 hpf when HSPCs are undergoing expansion, while *elf2b* is highly expressed. The low/absent expression of *elf1* may explain why it is not involved in HSPC expansion in zebrafish despite similar homology.

Here, we provide the first evidence that it is possible to pharmacologically enhance the clonal diversity of HSCs. Ro5-3335 induces transcriptional changes that increase RUNX1 binding to ELF factors, and this in turn promotes HSC stemness by activating cell cycle genes to encourage self-renewal and halt differentiation. Our color-based lineage tracing approach and engraftment assays did not show clonal dominance or lineage skewing of Ro5-3335 treated HSCs, but further investigation will be necessary to ensure that they do not eventually begin to expand or differentiate in a detrimental fashion that could lead to development of malignancy. Our work identifies a novel mechanism of action of Ro5-3335 and contributes to understanding the underlying mechanisms of HSC clonality. This may help to uncover new ways to enhance clonal diversity that could be very valuable for gene therapy and CRISPR targeting of blood stem cells in patients with blood disorders and malignancies. Ro5-3335 may even be a useful therapy for oligoclonal disorders such as bone marrow failures and aplastic anemia.

## Acknowledgements

The authors thank Nikolay Ogryzko and Stephen Renshaw for sharing the *mpeg1:NLSmClover* zebrafish line and John Bushweller for the CBFβ inhibitors. We would also like to thank Ran Jing, Chris Li and Michael Sanborn from George Daley’s lab for technical assistance. ALR received funding from an ALSF Young Investigator fellowship. LY received funding from a Croucher Fellowship for Postdoctoral Research. ARF, DFP and AMP were supported by the Cris Cancer Foundation Excellence Award (PR_EX_2020-24), the European Commission’s Horizon Europe program through the ERC Starting Grant MemOriStem (101042992), the Spanish National Research Agency (PID2020-114638RA-I00, PID2023-148073OB-I00), the EMBO Young Investigator Programme, and the CERCA Program/Generalitat de Catalunya. LIZ is a Howard Hughes Medical Institute Investigator. This work was supported by National Institute of Health grants R01HL144780-01, P01HL131477, U54DK110805, RC2DK12053, U01HL134812 and R24DK092760-05 (LIZ). We are exceptionally grateful to the Boston Children’s Hospital aquatics team and the BCH Flow Cytometry core. The authors would also like to acknowledge the technical assistance of the Functional Genomics Core Facility at IRB Barcelona (single-cell library preparations and sequencing).

## Author contributions

Conceptualization: ALR, LY, AC, LIZ; Methodology: ALR, LY, AC, CK, KDD; Formal Analysis: SY, LY, KDD, DFP, AMP; Investigation: ALR, LY, AC, DFP, AMP, CK, SJW, KDD, JM, SA, RB, RJF, VC, MCB; Visualization: ALR, LY, AC, CK; Funding acquisition: ALR, LY, AC, ARF, GQD, LIZ; Project administration: ALR, AC, JM; Supervision: ALR, GQD, ARF, LIZ; Writing – original draft: ALR, LY, AC, DFP, AMP; Writing – review & editing: ALR, LY, AC, CK, SW, KDD, SA, ARF, GQD, LIZ.

## Methods

### Animal models

Wild-type *casper* zebrafish^52^ and transgenic lines *Runx1+23:eGFP* (referred to as *Runx1:eGFP*)^37^, *mpeg1:NLSmClover* (referred to as *mpeg1:Clover*)^38^, c*myb:EGFP*^53^, *kdrl*:*Hsa.hras-mCherry* (referred to as *kdrl:mCherry*)^54^, *ubi:Zebrabow-M* (referred to as *ubi:Zebrabow*)^40^, *draculin:CreER^T2^* (referred to as *drl:CreER^T2^*)^55^ and *ubi:mCherry*^56^ were used in this study. Zebrabow insert number in *ubi:Zebrabow* zebrafish was quantified by qualitative PCR (qPCR) or droplet digital PCR (ddPCR) relative to single-insert zebrafish as described previously^6^, and zebrafish with ≥20 were chosen for breeding. All animals were housed at Boston Children’s Hospital, and all experiments and procedures were performed according to protocols approved by the Institutional Animal Care and Use Committee (IACUC) of Boston Children’s Hospital (Protocol Number 00002043).

### Zebrafish embryo culture screen

At 24 hours post fertilization (hpf), *Runx1:eGFP* embryos were dechorionated with Pronase (Sigma), washed with E3 embryo water, then resuspended in blastomere media composed of 85% LDF medium, 5% FBS, and 10% embryo extract. LDF medium contains 50% Leibowitz’s L-15 (Invitrogen), 20% DMEM (Invitrogen), and 30% DMEM/F-12 (Invitrogen), supplemented with 2% B27 (Gibco), 15 mM HEPES (Gibco), 1% L-glutamine (Gibco), 1% N2 (Gibco), 10 nM sodium selenite (Sigma), 0.018% sodium bicarbonate (Gibco), 0.04% Primocin (Invivogen), and 0.2% Penicillin-Streptomycin (Gibco). Following mechanical homogenization and filtering through a 40 μm nylon mesh filter, single cells were aliquoted into 384-well plates coated with 0.1% gelatin at 40 μl per well, resulting in approximately 2 embryo equivalents per well. Plates were immediately screened with chemicals from NIH, Library of Pharmacologically Active Compounds (Sigma), ICCB Known Bioactives (Biomol), Nuclear Hormone Receptor and Kinacore (ChemBridge) libraries at 30 μM. Cells were cultured in a 28°C incubator with 5% CO_2_ for 2 days. Cells were stained with Draq5 (Cell Signaling Technology) and imaged using a Cell Voyager 7000 (Yokogawa). Each chemical library plate was screened in duplicate. For analysis, a 4x image of the nuclear and fluorescent expression from each entire well was thresholded and the percent area was computed using ImageJ/Fiji. Control wells (200 or more per plate) were identified using quartile exclusion of outliers, and using these wells, a standard curve was built with GFP vs. nuclear staining in MatLab. From the standard curve, residuals were calculated for each treated well and divided by the standard deviation in the control wells to obtain the z-score of each chemical treatment.

### Drug treatment of zebrafish embryos

Clutch-matched, transgenic embryos were treated in 6-well plates at the indicated time points with 5 μM Ro5-3335 (Tocris) or DMSO (Sigma) as a vehicle control, diluted in E3 (n>10 embryos per treatment group). Plates were stored at 28.5°C until the relevant experimental time point, when live embryos were embedded in 0.8% low-melting point agarose containing Tricaine (0.16 mg/mL) in glass bottom 6-well plates and covered with E3 media containing Tricaine (0.16 mg/ml). For experiments involving time-lapse microscopy, 5 μM Ro5-3335 or DMSO was added to both the agarose and covering media to ensure consistent exposure to the drug. For transient treatment experiments in *mpeg1:Clover* embryos, as much media as possible was removed from wells and embryos were washed 4 times with fresh E3 media to ensure adequate removal of residual Ro5-3335, before being returned to fresh E3 media. The CBFβ–RUNX1 protein-protein interaction inhibitors, AI-14-91 and AI-10-104, and the corresponding control compound, AI-4-88, were received as a kind gift from the Bushweller Lab (Department of Molecular Physiology and Biological Physics, University of Virginia, Charlottesville, VA) and drug treatment assays were performed using the same methods as for Ro5-3335.

### Microscopy and image analysis

All single time-point and time-lapse microscopy was performed using a Yokogawa CSU-X1 spinning disk mounted on an inverted Nikon Eclipse Ti microscope equipped with dual Andor iXon EMCCD cameras and a climate controlled, motorized x-y stage. When necessary, screening of transgenes was performed using a Nikon SMZ18 stereomicroscope equipped with a Nikon DS-Ri2 camera. Animals were only included for imaging and analysis if expression of all transgenes could be identified. All images were acquired using NIS-Elements (Nikon), blinded, and processed using Imaris (Bitplane).

### Zebrafish morpholino injections

Morpholinos (GeneTools) were resuspended to 1 mM in nuclease free water, heated to 65°C for 5 minutes, and kept at room temperature. *Runx1:eGFP* embryos were injected into the yolk at the 1-4 cell stage with 0.125–0.25 pmol of morpholino, depending on toxicity. Morpholinos used in this study are listed in Supplementary Table 2.

### Zebrabow color labeling and analysis

Embryos generated from crossing *ubi:Zebrabow* and *drl:CreER^T2^* fish were treated with DMSO or Ro5-3335 at 24 hpf following our standard embryo drug treatment protocol, with the addition of 15 μM 4-hydroxytamoxifen (4-OHT). Plates were kept in the dark and then embryos were washed as described above for transient treatment experiments. Embryos with dim transgene expression were excluded, and the remaining embryos were raised to adulthood for analysis at 3-4 months. Kidney marrow was dissected under a Nikon SMZ1270 light microscope and mechanically dissociated by pipetting in cold 0.9x DPBS (Gibco) with 2% fetal bovine serum (FBS, Gemini Bio-Products) and 1 USP units/mL heparin (Sigma). Cell suspensions were filtered through a 40 μm filter and analyzed using a FACS Aria cell sorter (BD Biosciences). Color barcodes from marrow samples were quantified using previously published pipelines and adapted to a Python-based interface as previously described^6,7^. Granulocyte color output was selected as a read out of clonal changes, as these cells have a short half-life and reflect changes in the HSPC clonal output in a timely manner. Only zebrafish with greater than 75% recombination efficiency were processed. All samples were blinded prior to analysis and compared against clutch-matched siblings.

### Zebrafish HSPC transplantation and kidney marrow analysis

Double transgenic *Runx1:eGFP/ubi:mCherry* embryos were screened for mCherry expression then treated with DMSO or 5 μM Ro5-3335 from 24 hpf. At 72 hpf, embryos were finely chopped and dissociated using a razor blade followed by incubation in PBS containing Liberase (Roche) at 37°C for 20 minutes. Cell suspensions were filtered through a 40 μm filter and mCherry+/GFP+ cells were collected using a FACS Aria cell sorter (BD Biosciences). Collected cells were resuspended in PBS at an estimated concentration of 500 cells per microliter and 4 nanoliters were injected into the circulation of *casper* 2 dpf embryos via the Duct of Cuvier. Based on this dilution, the estimated cell number transplanted per embryo was 2 cells. At 3-4 months, whole kidney marrow (WKM) was analyzed for percentage of engrafted *Runx1:eGFP*+ cells using a LSR II flow cytometer (BD Biosciences). Recipients with GFP+ cells detected above background (>0.05% of WKM) were scored as engrafted, and expression of mCherry was analyzed to assess contribution to differentiated lineages. For secondary transplants, kidney marrow was collected from adult primary recipients (4 months old) by dissection under a Nikon SMZ1270 light microscope. Soft tissue was mechanically dissociated by pipetting in cold 0.9x DPBS (Gibco) with 2% fetal bovine serum (FBS, Gemini Bio-Products) and 1 USP units/mL heparin (Sigma). Cell suspensions were filtered through a 40 μm filter and GFP+ cells were collected using a FACS Aria cell sorter. Transplants were performed into 2 dpf *casper* embryos as described above, at an estimated number of 2 cells per embryo. Recipients were analyzed for percentage of engrafted *Runx1:eGFP*+ cells at 3 months using a LSR II flow cytometer.

### Zebrafish elf2b mutant generation

*Elf2b* sgRNA was synthesized from IDT (sequence in Supplementary Table 2). The gDNA was first assembled following the IDT manufacturer’s protocol by mixing with IDT Alt-R® CRISPR-Cas9 crRNA and IDT Alt-R® CRISPR-Cas9 tracrRNA and incubating at 95°C for 5 minutes in a thermocycler. The gRNA-Cas9 RNP was assembled by mixing 1 µL gDNA (6.1µM) with 1 µL Cas9 protein (1mg/mL, PNA Bio #CP01-50) and incubating at 95°C for 5 minutes in a thermocycler. The gRNA-Cas9 RNP injection mix was made by mixing 2 µL gRNA-Cas9 RNP, 2 µL H2O and 0.2 µL 50% Phenol Red. Each fish embryo was injected with 1 nL gRNA-Cas9 RNP injection mix at the one-cell stage. Mosaic F0 founders were grown to adulthood. Genomic DNA was extracted from fin clip tissue/embryos using the HotSHOT method. Briefly, genomic DNA was extracted with 100 µL 50 mM NaOH and incubated for 20 min at 95°C in a thermocycler, then neutralized with 10 µL 1 M Tris-HCl pH 7.5. The PCR was performed using Takara Advantage® 2 Polymerase (Takara #639202) following the manufacturer’s manual using *elf2b*_F and *elf2b*_R primers (sequences in Supplementary Table 2). Conditions for the PCR were as follows: 95°C, 5 min; [95°C, 30 sec; 68°C, 1 min] for 35 cycles; 68°C, 3 min; hold at 4°C. The amplicon was then purified using the DNA Clean Up kit (Zymo Research #D4033) and subjected to NGS or Sanger sequencing to identify the mutations. Germline-positive F0 founders were identified and crossed with wild type fish. The mutations of F1 embryos were analyzed using NGS or Sanger sequencing analysis.

### Zebrafish elf2b mutant maintenance and genotyping

The *elf2b*_7D mutant F1 fish line was identified to carry a frameshift mutation (7 bp deletion) and fish were maintained as heterozygotes. Embryos from heterozygous incrosses were used for the experiments. Single embryos were subjected to genotyping to identify wild-type, heterozygous and homozygous mutants using the NGS or Sanger sequencing analysis described above. Alternatively, for the *elf2b*_7D mutant identification, DNA gel electrophoresis following the *elf2b*_7D PCR was used for genotyping. Briefly, separate PCR reactions targeting the wild-type allele and the 7D mutant allele were done using Takara Advantage® 2 Polymerase (Takara #639202) following the manufacturer’s manual using different primers (sequences in Supplementary Table 2). Conditions for the PCR were as follows: 95°C, 5 min; [95°C, 30 sec; 68°C, 1 min] for 35 cycles; 68°C, 3 min; hold at 4°C.

### Hematopoietic stem cell isolation in mice

Following euthanasia, bone marrow was harvested from mouse femur, tibia, pelvis, and sternum through mechanical crushing, ensuring the retrieval of most HSCs. The collected bone marrow cells were then sieved through a 100 μm strainer and cleansed with a cold ‘Easy Sep’ buffer containing PBS with 2% FBS, followed by lysis of red blood cells using RBC lysis buffer (Biolegend, #420302). At first, mature lineage cells were selectively depleted through the Lineage Cell Depletion Kit, mouse (Miltenyi Biotec, #130-110-470), while the resulting Lin-(lineage-negative) fraction was then enriched for c-Kit expression using CD117 MicroBeads (Miltenyi Biotec, #130-091-224). These cKit-enriched cells were washed, blocked with FcX and stained with following fluorescently labeled antibodies: APC anti-mouse CD117, clone ACK2 (Biolegend, #105812), PE/Cy7 anti-mouse Ly6a (Sca-1) (Biolegend, #108114); Pacific Blue anti-mouse Lineage Cocktail (Biolegend, #133310); PE anti-mouse CD201 (EPCR) (Biolegend, #141504); PE/Cy5 anti-mouse CD150 (SLAM) (Biolegend, #115912); APC/Cyanine7 anti-mouse CD48 (Biolegend, #103432). Lentivirus production and HSC transduction were performed as described^57^.

### Mouse primary cell cultures

Ex-vivo cultures of HSCs were done under self-renewing F12-PVA-based conditions^58^. To this end, cell-culture activated 96-well flat-bottom plates were coated with a layer of 100 ng/mL fibronectin (Bovine Fibronectin Protein, CF R&D Systems #1030-FN) for 30 minutes at room temperature. Following the sorting process, HSCs were transferred into 200 μL of complete HSC media supplemented with 100 ng/mL recombinant mouse TPO and 10 ng/mL recombinant mouse SCF (PeproTech Recombinant Murine TPO, #315-14; PeproTech Recombinant Murine SCF, #250-03) and grown at 37°C with 5% CO2. During lentiviral library transduction, the first media change took place 24 hours post-transduction. 0.00025% DMSO (control) or 0.5 μM Ro5-5333 (treatment) was added to the culture media during the entire 14 days HSCs were maintained in the primary cultures. All other protocol steps followed previously published guidelines^58^.

### Single-cell encapsulation and library preparation for sequencing

For scRNA sequencing and subsequent plating, cells were pipetted up and down gently a few times to be dissociated into single cells and transferred to a 1.5 mL microtube. The well was then washed with prewarmed PBS to collect all the possible remaining cells. Cells were then concentrated by centrifugation at 800 g for 8 minutes. Washed cells were then blocked with FcX and stained with the E-SLAM stem cell antibodies panel, to confirm expansion of E-SLAM cells. Live cells were then sorted based on fluorescent reporter expression. Part of the sample as specified in text was then taken for constructing a single-cell library using Chromium Single Cell 3’ Reagent Kits (v3) following the manufacturer’s guidelines (10x Genomics). The remaining part was then plated back for further expansion in culture. To minimize the impact of batch effects on sequencing, we multiplexed different conditions leveraging the unique barcode pattern of our libraries together with Biolegend TotalSeq™ anti-mouse hashing antibodies, enabling the simultaneous preparation of libraries representing all experimental conditions in a single reaction for each day of sampling.

Following the reverse transcription of mRNA and first-strand cDNA amplification, 100 ng of the cDNA libraries were used as templates to amplify LARRY barcodes by nested PCR similar to the described methods^57^. The first PCR used forward primer (Pre-Enrichment forward) CTG AGC AAA GAC CCC AAC GAG AA together with the corresponding 10x Genomics dual index TruSeq reverse primer using the following conditions: 98°C, 3 min; [98°C, 20 sec; 58°C, 15 sec; 72°C, 20 sec] for 8 cycles; 72°C, 3 min; hold at 4°C. The PCR products were then purified with a 0.8:1 ratio of Ampure XP beads. Purified PCR products were then subjected to a second PCR using the forward primer (Trueseq_LARRY) GTG ACT GGA GTT CAG ACG TGT GCT CTT CCG ATC TGC TAG GAG AGA CCA TAT GGG ATC and the corresponding 10x dual index Truseq reserve primer, using the following conditions: 98°C, 3 min; [98°C, 20 sec; 58°C, 15 sec; 72°C, 20 sec] for 8 cycles; 72°C, 3 min; hold at 4°C. The final PCR products were then purified by a 0.8:1 ratio of Ampure XP bead: PCR products, were indexed using the 10x dual index TruSeq kit, and sequenced using Illumina NovaSeq or NextSeq.

### scRNA-seq data processing and calling of lineage barcodes

Generation of single-cell matrices for gene expression and LARRY lineage barcodes was performed using *cloneRanger*^41^ to process 10X Genomics single-cell RNA-seq together with LARRY barcoding using default parameters. A Seurat^59^ object containing single-cell count matrices from GEX, LARRY and TotalSeq™ counts was created with the function Read10x from Seurat. Finally, cell doublets were removed with scDblFinder using default parameters and TotalSeq™ sequences were demultiplexed with the function hashedDrops from the DropletUtils R package^60^ with default parameters. The assignment of LARRY barcodes to individual cells was performed by cloneRanger: first, we generated a filtered LARRY matrix by removing barcode UMIs that were sustained by less than 5 sequencing reads (this information is stored in the *molecule_info.h5* file generated by cellranger). Then, we further filtered the LARRY matrix by removing all barcodes with less than 3 UMIs. After filtering, barcodes were assigned to individual cells as following: (a) for cells in which only one barcode was detected after filtering, that barcode was assigned, (b) for cells in which more than one barcode was detected post-filtering, the top barcode with higher UMI counts was assigned and (c) for cells in which there were ties in the top barcode, no barcode was assigned. This barcode calling strategy was developed to minimize mixing cells from different clones at the expenses of having higher chances to split real clones into subclones.

### Single-cell integration, clustering and annotation

To integrate scRNA-seq samples, we applied the *IntegrateLayers* Seurat v5 workflow with the Reciprocal PCA algorithm to integrate the single-cell GEX matrices. We followed a standard Seurat pipeline with some minor modifications. Raw counts were normalized with the function *NormalizeData* and the top 3000 variable genes were selected. From those, we removed all ribosomal and mitochondrial genes, as well as genes that correlated with cell cycle genes (*Ube2c*, *Hmgb2*, *Hmgn2*, *Tuba1b*, *Ccnb1*, *Tubb5*, *Top2a*, *Tubb4b*, pearson correlation of 0.1 or more). Then, filtered variable genes were used to compute the top 50 reciprocal-PCA components. The kNN graph was computed using the function *FindNeighbors* setting the number of neighbors to 30. Clusters were generated with the function *FindClusters* with a resolution of 0.3 and the Louvain algorithm.

### Differential abundance testing

Differential abundance testing across conditions was performed using the miloR package^61^ (using default parameters unless otherwise indicated). In order to generate the Milo object and the kNN graph, the same number of Principal Components and Neighbours used to compute the UMAP were used for this analysis. The proportion of randomly sampled cells for the generation of neighborhoods was set to 0.1. Neighborhoods that had less than 70% of cells corresponding to a unique cell type were labeled as “Mixed”. Neighborhoods with FDR < 0.1 were considered as differentially enriched between DMSO and Ro5-5333 samples.

### Quantification and annotation of HSC clonal behaviors

Quantification of HSC clonal behaviors followed the published methodology^41^, relying on cellular distribution patterns between HSC and non-HSC clusters. Output activity (A*_i_*) was calculated as the ratio of the clone *i* frequency in non-HSC clusters to its frequency in the HSC cluster plus a pseudocount of 0.001.

### hiPSC derived hematopoietic progenitor culture and analysis

Hemogenic endothelial (HE) cells were differentiated from hiPSCs as previously described^62^. In brief, hiPSCs were cultured as embryoid bodies (EBs) for 8 days in 5% CO_2_/5%O_2_/90% N_2_. HE cells were enriched by magnetic-activated cell sorting (MACS; Miltenyi) for CD34+ cells from day 8 EBs and plated with indicated concentrations of Ro5-3335 or DMSO control in EHT medium (EB medium with BMP4 (10 ng/mL), bFGF (5 ng/mL), IL-3 (30 ng/mL), IL-6 (10 ng/mL), IL-11 (5 ng/mL), insulin-like growth factor 1 (25 ng/mL), vascular endothelial growth factor (5 ng/mL), stem cell factor (100 ng/mL), erythropoietin (2 U/mL), thrombopoietin (30 ng/mL), FMS-like tyrosine kinase 3 ligand (10 ng/mL), Sonic hedgehog (20 ng/mL), angiotensin II (10 μg/mL) and losartan (100 μM)). Half media addition was performed on day 3 or 4 of EHT. On day 4 of EHT both floating and adherent cells were assayed by flow cytometry for CD34 and CD45 expression, while on day 7 only floating cells were harvested and analyzed. Data acquisition and processing was performed on a BD LSRFortessa cell analyser and using FlowJo 10.4.2 software (Tree Star). Antibodies used were Mouse anti-Human CD34 PE-Cy7, Clone: 581 (Fisher Scientific) and Mouse anti-Human CD45 APC, Clone: J33 (Beckman Coulter), both at 1:100 dilutions.

### CD34+ cell culture and drug treatment

Human CD34+ cells, isolated from peripheral blood of granulocyte colony-stimulating factor-mobilized healthy volunteers, were purchased from the Fred Hutchinson Cancer Research Center. Cells from more than 5 different donors were used in this study. The cells were maintained and differentiated as previously described^63,64^. At day 0, cells were thawed and plated down in expansion media (StemSpan™ SFEM II (Stem Cell Technologies Inc, #09655), 1% StemSpan™ CC100 (Stem Cell Technologies Inc, #2690), 1% Penicillin-Streptomycin (5,000 U/mL, Gibco, #15070063). Cells were recovered for approximately 6 hours before drug treatment. For chemical treatment, cells were plated at a density of 80 K/mL. Ro5-3335 (5 µM) or the equivalent volume of DMSO was added at day 0. At day 4, an equal volume of fresh expansion media with compounds was supplemented. At day 6, cells were quantified and prepared for FACS analysis. Briefly, cells were washed with FACS buffer (PBS, 1% FBS) and pelleted down at 300g for 5 minutes. Cell pellets were gently resuspended in 100 µL FACS buffer and stained with antibodies (CD38-APC, Clone: HIT2 (Biolegend, 1:100), CD45RA-PE-Cy7, Clone: HI100 (Biolegend, 1:100), CD34-FITC, Clone: 581 (Biolegend, 1:50), and CD90-PE Clone: 5E10 (Biolegend, 1:33)) at 4°C for 30 minutes. After antibody staining, cells were washed with FACS buffer and pelleted down at 300g for 5 minutes. Cell pellets were gently resuspended in 300 µL FACS buffer and filtered through FACS filter tubes with a 35 µm nylon mesh. FACS sorters (Sony MA900 cell sorter and BD Aria III) were carefully calibrated, and compensation was performed before analysis. Before running each sample, Sytox Blue (1:100, ThermoFisher, # S34857) was added to the cell suspension for live/dead gating. For expansion analysis, cells were quantified at day 0 and a portion of cells were subjected to FACS analysis. To identify the most immediate epigenetic changes in response to Ro5-3335 treatment, cells were stimulated for 24 hours with 20 µM Ro5-3335 after 6 days of expansion and harvested for ChIP-seq, CUT&RUN and ATAC-seq assays.

### EdU incorporation and staining assay

EdU assays were performed following the Click-iT™ Plus EdU Alexa Fluor™ 647 Flow Cytometry Assay Kit (ThermoFisher, C10634) manufacturer’s guidelines. Briefly, 10 µM EdU was added to cells and incubated for 1 hour before harvest. Before fixation, cells were stained with cell surface marker antibodies (CD38-V450 Clone: HIT2 (Biolegend, 1:100), CD45RA-PE-Cy7, Clone: HI100 (Biolegend, 1:100), CD34-FITC, Clone: 581 (Biolegend, 1:50), and CD90-PE, Clone: 5E10 (Biolegend, 1:33)) as previously described. After antibody staining, cells were fixed, permeabilized and stained with Alexa-Flour 647 Azide.

### Chromatin Immunoprecipitation (ChIP)

For ChIP-seq experiments, the following antibodies were used: RUNX1 (Abcam ab23980), CBFβ (Abcam ab195411), ELF1 (Santa Cruz sc133096X), and H3K27ac (Abcam ab4729). ChIP experiments were performed as previously described with slight modifications^64,65^. Briefly, 20 - 30 million cells for each ChIP were crosslinked by the addition of 1/10 volume 11% fresh formaldehyde for 10 minutes at room temperature. The crosslinking was quenched by the addition of 1/20 volume 2.5 M Glycine. Cells were washed twice with ice-cold PBS and the pellet was flash-frozen in liquid nitrogen. Cells were kept at −80°C until the experiments were performed. Cells were lysed in 10 mL of Lysis buffer 1 (50 mM HEPES-KOH, pH 7.5, 140 mM NaCl, 1 mM EDTA, 10% glycerol, 0.5% NP-40, 0.25% Triton X-100, and protease inhibitors) for 10 minutes at 4°C. After centrifugation, cells were resuspended in 10 mL of Lysis buffer 2 (10 mM Tris-HCl, pH 8.0, 200 mM NaCl, 1 mM EDTA, 0.5 mM EGTA, and protease inhibitors) for 10 minutes at room temperature. Cells were pelleted and resuspended in 1 mL of Sonication buffer (10 mM Tris-HCl, pH 8.0, 100 mM NaCl, 1 mM EDTA, 0.5 mM EGTA, 0.1% Na-Deoxycholate, 0.05% N-lauroylsarcosine, and protease Inhibitors) and sonicated in a Bioruptor sonicator for 24-40 cycles of 30s followed by 1 min resting intervals. Samples were centrifuged for 10 min at 18,000 g and 1% of TritonX was added to the supernatant. Prior to the immunoprecipitation, 50 mL of protein G beads (Invitrogen #100-04D) for each reaction were washed twice with PBS, 0.5% BSA twice. Finally, the beads were resuspended in 250 mL of PBS, 0.5% BSA and 5 mg of each antibody. Beads were rotated for at least 6 hours at 4°C and then washed twice with PBS, 0.5% BSA. Cell lysates were added to the beads and incubated at 4°C overnight. Beads were washed 1x with (20 mM Tris-HCl (pH 8), 150 mM NaCl, 2mM EDTA, 0.1% SDS, 1%Triton X-100), 1x with (20 mM Tris-HCl (pH 8), 500 mM NaCl, 2 mM EDTA, 0.1% SDS, 1%Triton X-100), 1x with (10 mM Tris-HCl (pH 8), 250 nM LiCl, 2 mM EDTA, 1% NP40) and 1x with TE and finally resuspended in 200 mL elution buffer (50 mM Tris-Hcl, pH 8.0, 10 mM EDTA and 0.5%–1% SDS). Prior to addition to the beads, 50 µL of cell lysates were kept as input. Crosslinking was reversed by incubating samples at 65°C for at least 6 hours. Afterwards the cells were treated with RNase and proteinase K and the DNA was extracted by Phenol/Chloroform extraction.

### RNA sequencing (RNA-seq)

RNA from 1 million cells was isolated using Trizol according to the manufacturer’s instructions. The RNA was DNAse-treated using the RNase-free DNase set from Qiagen (#79254) according to the manufacturer’s instructions. Total RNA was treated with the Ribo-Zero Gold kit (Human/Mouse/Rat, Epicentre) according to the manufacturer’s instructions. Briefly, 225 μL of magnetic beads per sample were washed in RNAse-free water five times. After the last wash, 65 μL of Magnetic Bead resuspension solution was added and the beads were kept at room temperature (RT). For each sample, the recommended amount was used according to the manufacturer and the recommended reaction was set-up and incubated at RT for 5 minutes. The mixture was then transferred to the magnetic beads and incubated at RT for 5 minutes and 50°C for 5 minutes. The Ribo-Zero treated RNA was then purified using the recommended modified protocol for RNeasy MinElute Cleanup Kit. Finally, the Ribo-Zero treated RNA was used to create multiplexed RNA-seq libraries using the ScriptSeq™ v2 RNA-Seq Library Preparation Kit (Epicentre) according to the manufacturer’s instructions. Briefly 500 pg of Ribo-Zero treated RNA was fragmented and used to produce cDNA according to the manufacturer’s protocol. The cDNA was cleaned with Agencourt AMPure purification kit and this was used as a template to produce multiplexed libraries (see below).

### ChIP-seq and RNA-seq library preparation

Briefly, ChIP-seq libraries were prepared using the following protocol. End repair of immunoprecipitated DNA was performed using the End-It End-Repair kit (Epicentre, ER81050) and incubating the samples at 25°C for 45 minutes. End-repaired DNA was purified using AMPure XP Beads at 1.8x the reaction volume (Agencourt AMPure XP – PCR purification Beads, BeckmanCoulter, A63881) and separating beads using DynaMag-96 Side Skirted Magnet (Life Technologies, #12027). A-tail was added to the end-repaired DNA using NEB Klenow Fragment Enzyme (3’-5’ exo, M0212L), 1x NEB buffer 2 and 0.2 mM dATP (Invitrogen, #18252-015) and incubating the reaction mix at 37°C for 30 minutes. A-tailed DNA was cleaned up using AMPure beads (1.8x of reaction volume). Subsequently, cleaned up A-tailed DNA went through Adaptor ligation reaction using Quick Ligation Kit (NEB, M2200L) following the manufacturer’s protocol. Adaptor-ligated DNA was first cleaned up using AMPure beads (1.8x of reaction volume), eluted in 100 μL and then size-selected using AMPure beads (0.9x the final supernatant volume, 90 μL). Adaptor-ligated DNA fragments of proper size were enriched with PCR reaction using Phusion High-Fidelity PCR Master Mix kit (NEB, M0531S) and specific index primers supplied in NEBNext Multiplex Oligo Kit for Illumina (Index Primer Set 1, NEB, E7335L). Conditions for the PCR were as follows: 98°C, 30 sec; [98°C, 10 sec; 65°C, 30 sec; 72°C, 30 sec] for 15 to 18 cycles; 72°C, 5 min; hold at 4°C. PCR enriched fragments were further size-selected by running the PCR reaction mix in 2% low-molecular weight agarose gel (Bio-Rad, #161-3107) and subsequently purifying them using QIAquick Gel Extraction Kit (28704). Libraries were eluted in 25 μL elution buffer. After measuring concentration using a Qubit, all the libraries went through quality control analysis using an Agilent Bioanalyzer. Samples with proper size (250-300 bp) were selected for Next-Generation Sequencing using the Illumina HiSeq 2000 or 2500 platform.

For the RNA-seq libraries, purified double-stranded cDNA underwent end-repair and dA-tailing reactions following manufacturer’s reagents and reaction conditions. The obtained DNAs were used for Adaptor Ligation using adaptors and enzymes provided in NEBNext Multiplex Oligos for Illumina (NEB#E7335) and following the kit’s reaction conditions. Size selection was performed using AMPure XP Beads (starting with 0.6x the reaction volume). DNA was eluted in 23 μL of nuclease free water. Eluted DNA was enriched by PCR using Phusion High-Fidelity PCR Master Mix kit (NEB, M0531S) and specific index primers supplied in NEBNext Multiplex Oligo Kit for Illumina (Index Primer Set 1, NEB, E7335L). Conditions for the PCR were as follows: 98°C, 30 sec; [98°C, 10 sec; 65°C, 30 sec; 72°C, 30 sec] for 15 cycles; 72°C, 5 min; hold at 4°C. PCR reaction mix was purified using Agencourt AMPure XP Beads and eluted in a final volume of 20 μL. After measuring the concentrations using a Qubit, all the libraries went through quality control analysis using an Agilent Bioanalyzer. Samples with proper size (250-300 bp) were selected for High-Throughput Sequencing using the Illumina HiSeq 2500 platform.

### ChIP-seq and CUT& RUN data analysis

All ChIP-Seq and CUT&RUN datasets were aligned to UCSC build version hg19 of the human genome using Bowtie2 (version 2.2.1)^66^ with the following parameters: --end-to-end, -N0, -L20. We used the MACS2 version 2.1.0^67^ peak finding algorithm to identify regions of ChIP-seq and CUT&RUN peaks, with a q-value threshold of enrichment of 0.05 for all datasets.

### RNA-seq data analysis

Quality control of RNA-seq datasets was performed by FastQC (Babraham Bioinformatics) and Cutadapt^68^ to remove adaptor sequences and low quality regions. The high-quality reads were aligned to UCSC build hg19 of human genome using Tophat 2.0.11^69^ without novel splicing form calls. Transcript abundance and differential expression were calculated with Cufflinks 2.2.1^70^. FPKM values were used to normalize and quantify each transcript, and the resulting list of differentially expressed genes were filtered by log fold change > 1 and q-value > 0.05.

### Assay for Transposase Accessible Chromatin (ATAC-seq)

CD34+ cells were expanded and differentiated using the protocol mentioned above. 5×10^4^ cells per differentiation stage were harvested by spinning at 500 x g for 5 minutes, at 4°C. Cells were washed once with 50 μL of cold PBS and spun down at 500 x g for 5 minutes at 4°C^71^. After discarding supernatant, cells were lysed using 50 μL cold lysis buffer (10 mM Tris-HCl pH 7.4, 10 mM NaCl, 3 mM MgCl2, 0.1% IGEPAL CA-360) and spun down immediately at 500 x g for 10 minutes at 4°C. Cells were then precipitated, kept on ice, and subsequently resuspended in 25 μL 2x TD Buffer (Illumina Nextera kit), 2.5 μL Transposase enzyme (Illumina Nextera kit, 15028252) and 22.5 μL nuclease-free water in a total of 50 μL per reaction for 1 hour at 37°C. DNA was then purified using the Qiagen MinElute PCR purification kit (#28004) in a final volume of 10 μL. Libraries were constructed according to the Illumina protocol using the DNA treated with transposase, NEB PCR master mix, SYBR^TM^ green, universal and library-specific Nextera index primers. The first round of PCR was performed under the following conditions: 72°C, 5 min; 98°C, 30 sec; [98°C, 10 sec; 63°C, 30 sec; 72°C, 1 min] x 5 cycles; hold at 4°C. Reactions were kept on ice and using a 5 μL reaction aliquot, the appropriate number of additional cycles required for further amplification was determined in a side qPCR reaction: 98°C, 30 sec; [98°C, 10 sec; 63°C, 30 sec; 72°C, 1 min] x 20 cycles; hold at 4°C. Upon determining the additional number of PCR cycles required for each sample, library amplification was conducted using the following conditions: 98°C, 30 sec; [98°C,10 sec; 63°C, 30 sec; 72°C, 1 min] x appropriate number of cycles; hold at 4°C. Libraries prepared went through quality control analysis using an Agilent Bioanalyzer. Samples with appropriate nucleosomal laddering profiles were selected for Next Generation Sequencing using the Illumina HiSeq 2500 platform.

### ATAC-seq data analysis

All human ATAC-seq datasets were aligned to build version hg19 of the human genome using Bowtie2 (v2.2.1)^66^ with the following parameters: --end-to-end, -N0, -L20. Coverage files for display were created using MACS with parameters -w -S –space=50 –nomodel –shiftsize=200. We used the MACS2 version 2.1.0 peak-finding algorithm^67^ to identify regions of ATAC-seq peaks, with the following parameters --nomodel --shift −100 --extsize 200. A q-value threshold of enrichment of 0.05 was used for all datasets. For correlation of ATAC-seq data with ChIP-seq binding, reads were mapped to the human genome (hg19) using Bowtie (v2.2.5)^66^ with default options. Read counts were normalized by library size to get counts per million (CPM).

### Transcription factor motif identification at DNA regions with increased binding of RUNX1 and increased gene expression upon Ro5-3335 treatment

The motifs enriched in transcription factor (TF) binding sites represented by ChIP-seq peaks, or enhancers represented by ATAC-seq peaks, were analyzed using the findMotifsGenome.pl program in the HOMER package (UCSD), as previously described^72^. For the motif search, background sequences used were either random human genome sequences or corresponding peaks of interest. The most significant known Homer motifs and *de novo* motifs by P-value were calculated. The transcription factor occupancy profile plots were generated using the deepTools2 suite^73^ with the computeMatrix and plotHeatmap commands in the reference-point mode, where the TF binding profiles were plotted over the 2kb upstream and downstream regions of every TF binding site. The heatmaps are sorted by the total intensity of each region.

### Comparison of Ro5-3335-mediated gene expression at RUNX1 and ELF1 co-bound vs RUNX1 only DNA regions

For these analyses, we assigned each of the 43,628 RUNX1 ChIP-seq peaks and 10,885 ELF1 peaks to their closest genes by adjacency using the UROPA package, using 100kb around the TSS of a gene as a criterion. Next, for genes that were assigned with RUNX1 or ELF1 binding sites, there were 4,791 genes containing RUNX1 and ELF1 co-binding regions, and 1,243 genes containing both RUNX1 and ELF1 individual binding sites but no co-binding. There were 4,151 genes containing only RUNX1 binding sites and 1,971 genes containing only ELF1 binding sites. We considered a peak intensity increase as a larger than 2-fold increase in RUNX1 binding. 2,890 genes contained increased RUNX1-ELF1 co-binding peaks, and 3,236 genes contained increased RUNX1-only peaks.

### Western blotting

Antibodies used for co-immunoprecipitation and ChIP-seq studies were tested using western blot analysis. For this purpose, cells were treated for 2 hours, washed in 1x PBS, and collected in RIPA buffer with protease and phosphatase inhibitors. Samples were run on acrylamide gel and transferred onto a nitrocellulose membrane. Membrane was blocked for one hour in 5% milk or TBS-T and incubated at RT with anti-Runx1 (Santa Cruz SC-365644X), anti-ELF1 (Santa Cruz SC-133096X) or anti-ELF4 (Santa Cruz SC-390689X). The membranes were washed, incubated with HRP-conjugated secondary antibodies for 1 hour at RT, and developed with SuperSignal West Pico Plus Chemiluminescent substrate.

### Co-immunoprecipitation (Co-IP)

Co-IP was performed as previously described^74^. Cells were washed twice with 1x PBS. Nuclei were isolated with 0.05% Triton in PBS and lysed in nuclei lysis buffer (20mM Hepes-KOH pH7.9, 25% glycerol, 420mM NaCl, 1.5mM MgCl2, 0.2mM EDTA, 0.5mM DTT). 500 μg of protein extract was used and the salt concentration was diluted from 420mM to 150mM NaCl using 20mM Hepes-KOH pH7.9, 20% glycerol, 0.25 mM EDTA, 0.05% NP-40. Cell lysates were pre-cleared with non-antibody bounds beads for 1 hour at 4°C. 5 mg antibody-bound protein G Dynabeads were added to pre-cleared lysate and samples incubated overnight at 4°C. Antibodies used were IgG (Cell Signalling #2729) and anti-Runx1 (Santa Cruz SC-365644X). Protein-bead complexes were then washed 5 times with wash buffer (20 mM Hepes-KOH pH7.9, 10% Glycerol, 150 mM NaCl, 1.5 mM MgCl2, 0.2 mM EDTA, 0.5 mM DTT) and beads were boiled in 50 μL Laemmli Buffer for 15 minutes at 95°C to elute proteins. Subsequently, samples were subjected to western blot as described above.

### CUT&RUN assay

Human mobilized PB CD34+ HSPCs were purchased from the Fred Hutchinson Cancer Research Center. The cells were maintained and differentiated as previously described^63,64^. On day 0, cells were thawed and plated in expansion media (StemSpan, CC100, 1:100, 1% Pen/Strep). On day 6, cells were treated with 20 µM Ro5-3335 for 24 h. On day 7, cells were harvested and washed 3 times with ice-cold PBS. Cells were pelleted at 600 g for 3 min at RT, then subjected to the CUT&RUN assay following the manufacturers protocol (CUTANA, CUT&RUN Kit Version 6). Briefly, 500,000 cells per condition were resuspended in Wash Buffer (Pre-Wash Buffer, 25X Protease Inhibitor, 1 M Spermidine, 1000x Wash Buffer Enhancer 1, 2500x Wash Buffer Enhancer 2) then spun twice at 600g for 3 min at RT. Activated ConA beads were added to washed cells, gently vortexed then incubated for 10 min at RT to adsorb cells to beads. Tubes were placed on a magnet then the slurry was resuspended in cold Antibody Buffer (Wash Buffer, 5% Digitonin, 0.5 M EDTA) prior to the addition of 0.5 μg of primary antibody to each experimental reaction (RUNX1, Abcam AB23980; ELF1, EpiCypher #13-2023; ELF2, Proteintech #12499-1-AP) or IgG control antibody (CUTANA). Cells were incubated overnight on a nutator at 4°C. The next day, tubes were placed on a magnet and the slurry was resuspended in cold Cell Perm Buffer (Wash Buffer, 5% Digitonin) 3 times. After the addition of pAG-Mnase, cells were incubated for 10 min at RT then returned to the magnet and resuspended in cold Cell Perm Buffer 2 times. Cells were incubated on a nutator for 2 hr and 4°C after the addition of 100 mM Calcium Chloride. At the end of the incubation, Stop Master Mix was added to each reaction (E. coli Spike-in DNA, Stop Buffer) then tubes were placed in a thermocycler at 37°C for 10 min. Tubes were returned to the magnet then the supernatants containing CUT&RUN-enriched chromatin were transferred to new tubes and CUTANA DNA Purification Beads were added. Following at 5 min incubation at RT, tubes were placed on the magnet for 2-5 min then resuspended in 85% EtOH two times. Beads were air dried for 2-3 min at RT then the DNA was eluted in 0.1X TE Buffer.

### CUT&RUN library preparation

5 ng CUT&RUN DNA was used as input for CUTANA CUT&RUN Library Prep (# 14-1001), which was carried out following manufacturer’s guidelines. Briefly, DNA was end-repaired followed by adapter ligation and U-excision. Modified DNA was purified using SPRIselect bead selection and eluted in 0.1x TE. Samples were indexed with unique i7/i5 primer combinations and PCR amplified using 14 cycles. Resulting libraries were purified using SPRIselect bead selection and eluted in 0.1x TE. Library size and integrity was evaluated using Agilent Tapestation with D1000 ScreenTape, per manufacturer’s recommendations, to confirm peak enrichment from 200-700 bp. Samples were then pooled at equimolar ratios and sequenced at 2×150 bp paired-end reads with 360 million total read depth via Illumina sequencing with Quintara Biosciences.

### Statistical analysis

The detailed methodologies used for statistical tests and the resulting significance values obtained comparing the control and test groups are described in the relevant figure legends, figures and method sections. Biological replicates and observed data-point variations are mentioned wherever applicable. All statistical analyses were carried out using the statistical computing/graphics software R, GraphPad Prism and Excel (Microsoft). Error bars indicate mean +/− SEM or +/− SD, as denoted in figure legends. Unless specified, *P-*values were obtained with two-tailed Student’s *t-*test for comparisons between two sample groups, or One-way ANOVA for comparisons between multiple groups, as denoted in figure legends. For zebrafish experiments, siblings were used to avoid bias from transgene variability, then randomized into vehicle control and Ro5-3335 treatment conditions. For mouse and human cell experiments, cells from the same batch/donor were split and randomized into vehicle control and Ro5-3335 treatment conditions.

