## Supplemental materials for "A molecular switch: Ro5-3335 drives hematopoietic stem cell division and clonality through distinct RUNX1 binding partners"

### Supplementary Figure 1

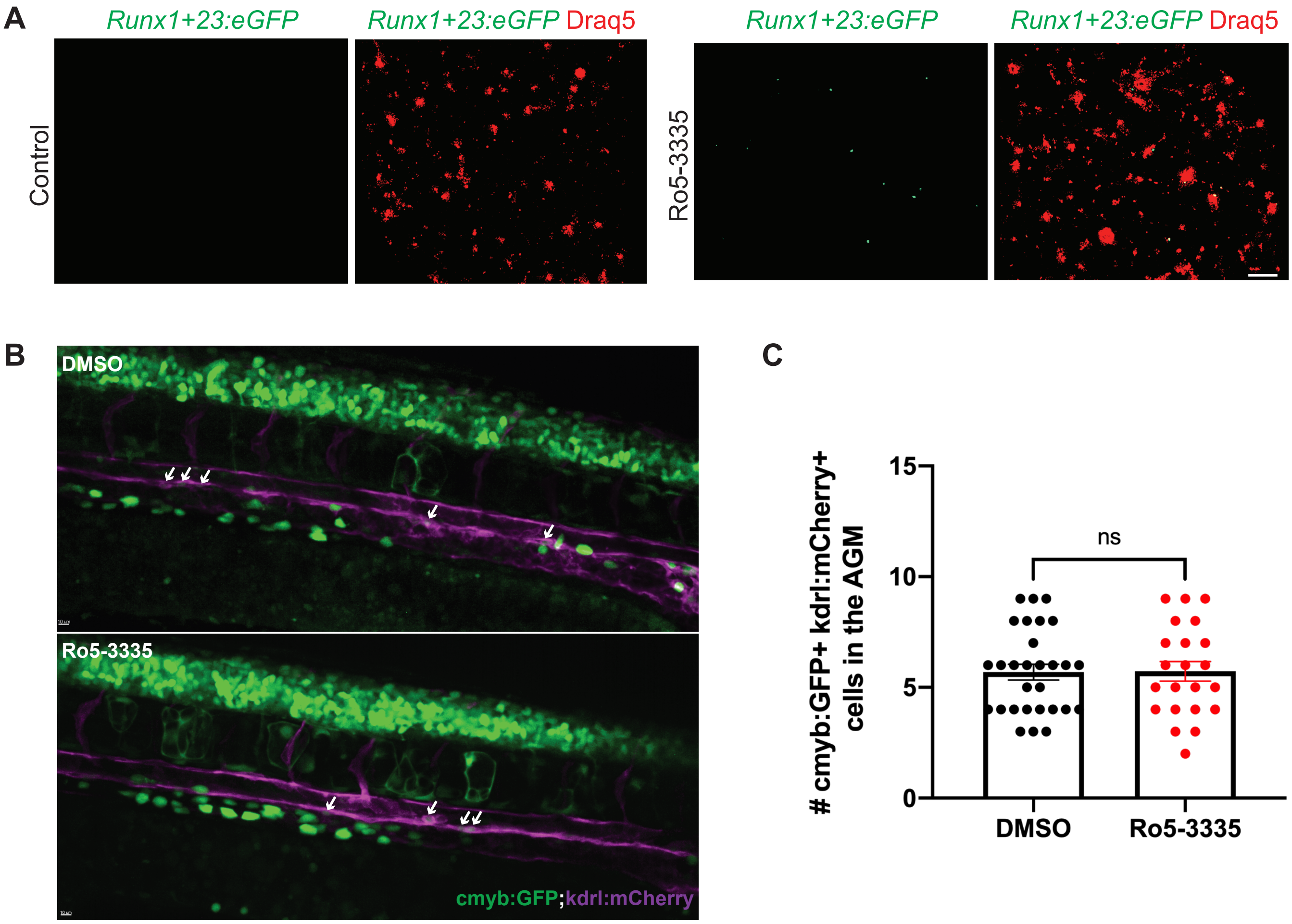

Supplementary Figure 2

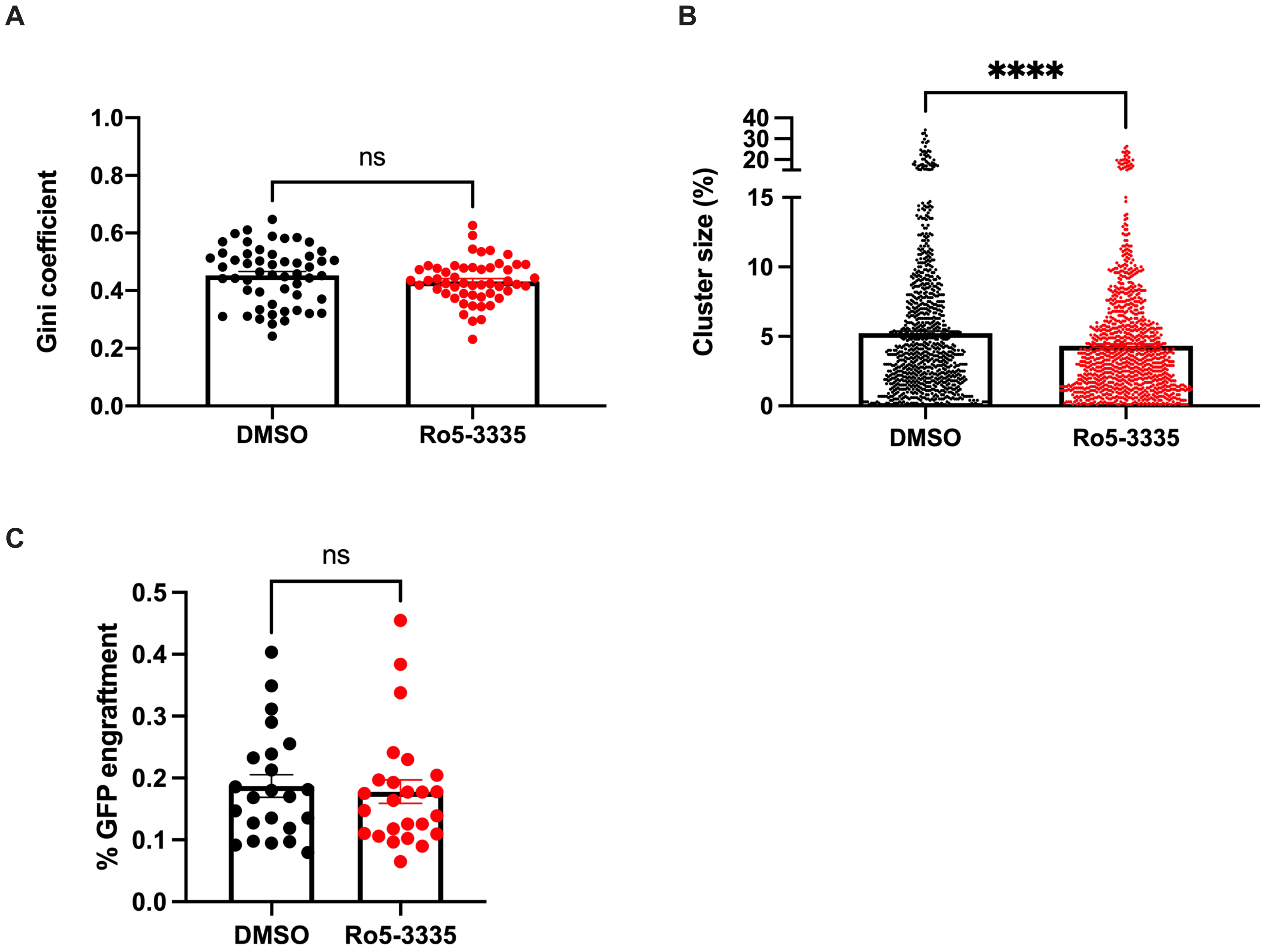

Supplementary Figure 3

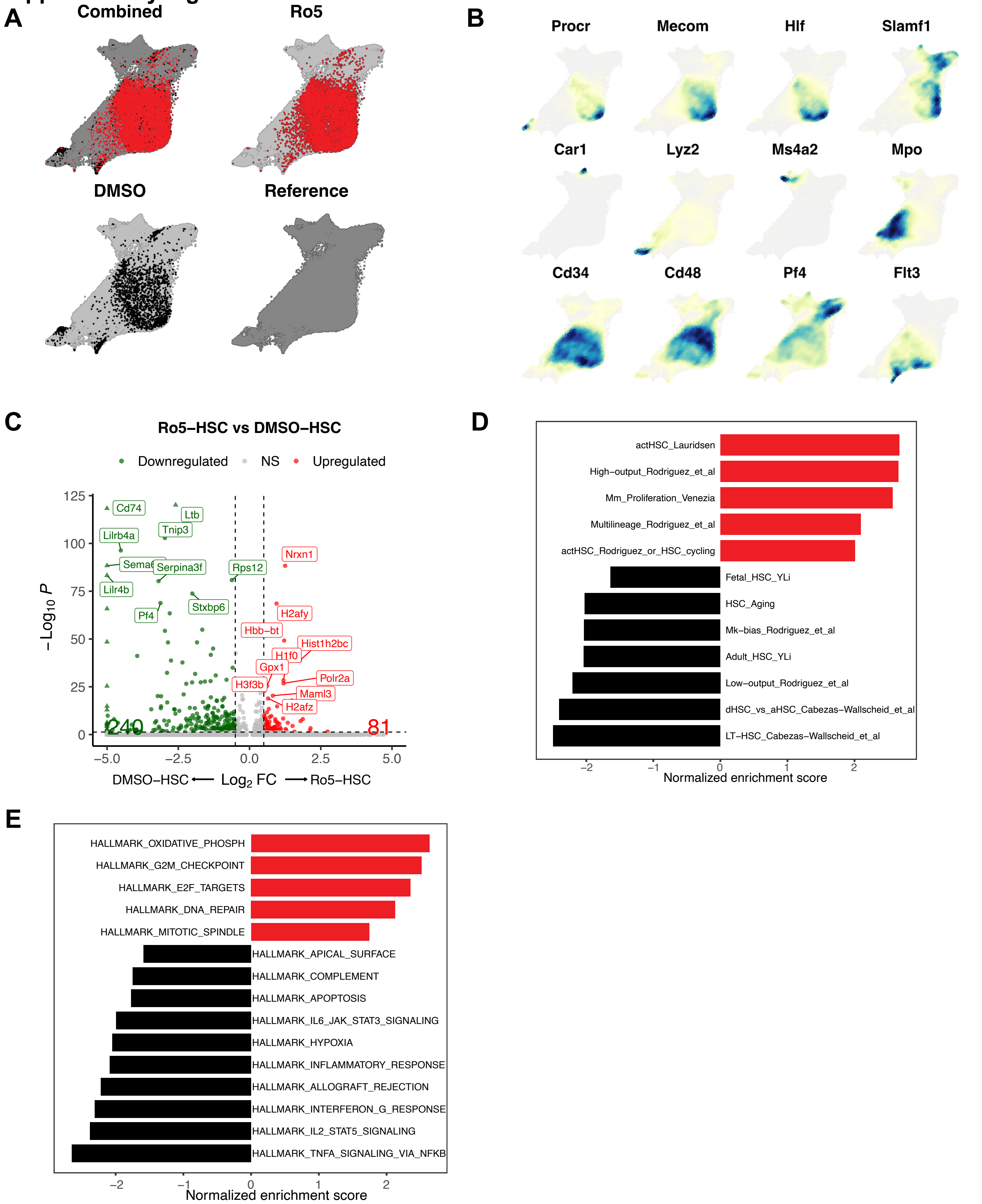

### Supplementary Figure 4

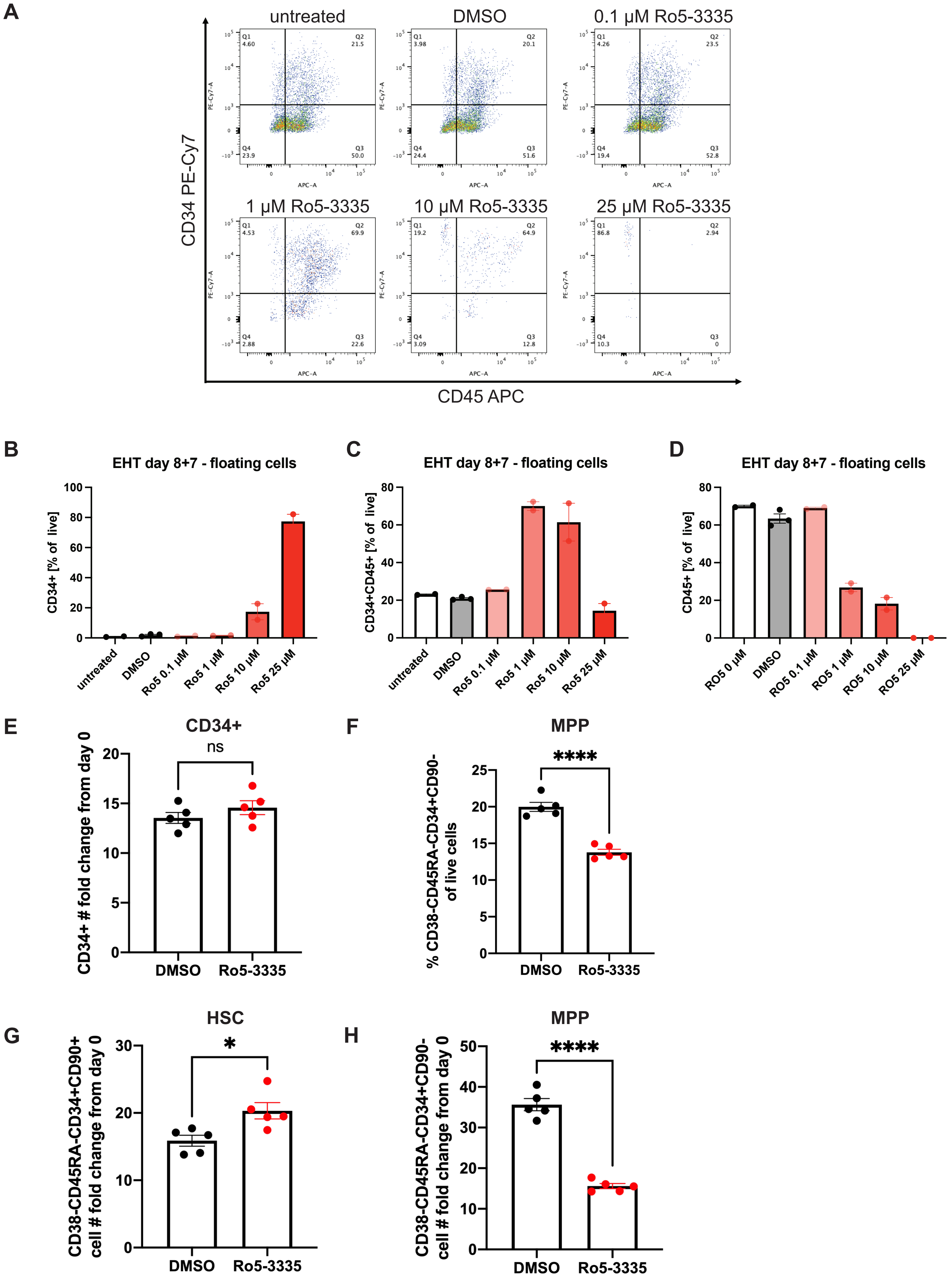

### Supplementary Figure 5

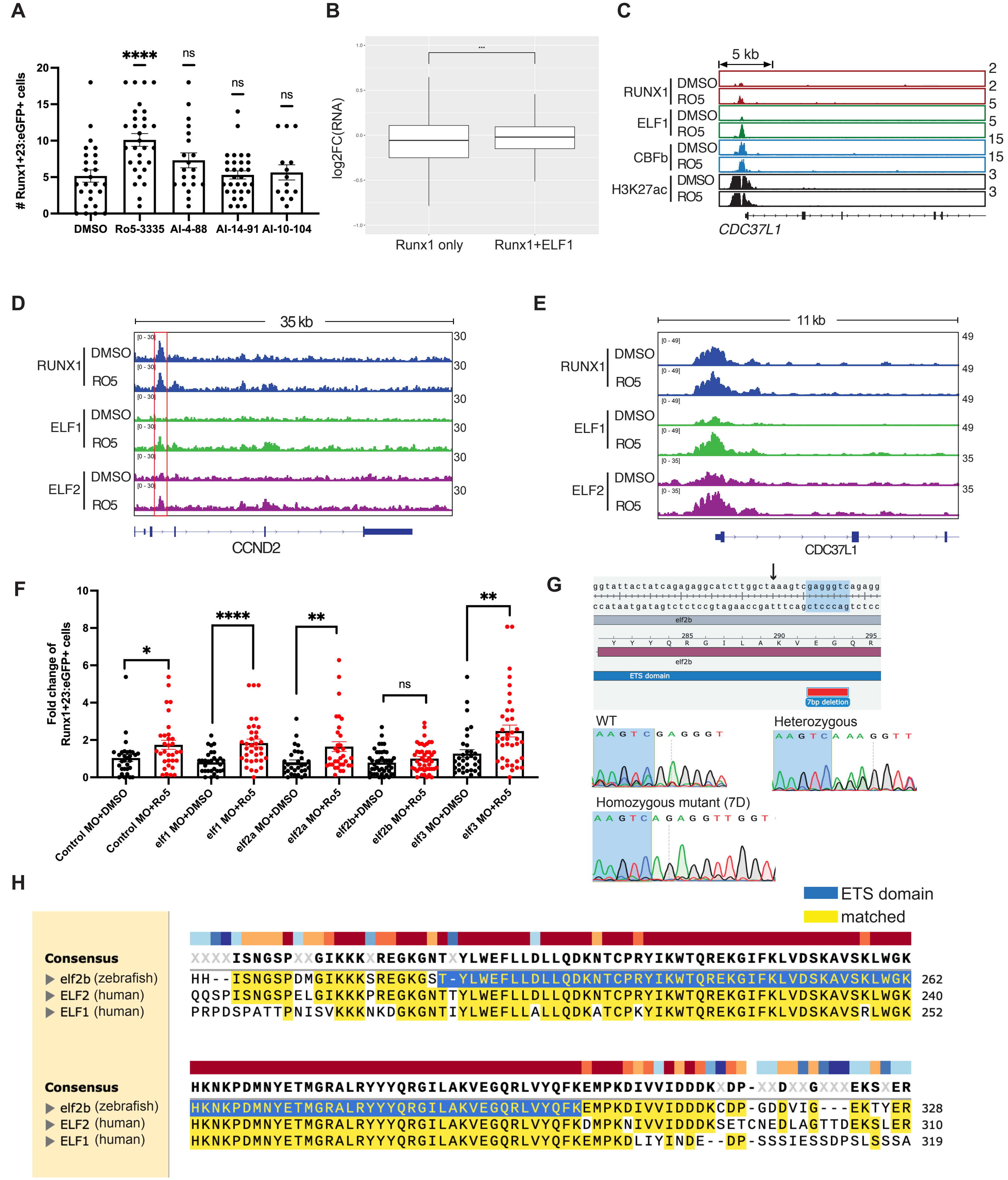

#### Supplementary Table 1

| Table S1 - Chemical modulators of Runx1:GFP in zebrafish embryo cultures. |
|---|
| Ro5-3335 |
| BI-D1870 |
| 5-methyl-2-(3-([methyl(pyridin-2-ylmethyl)amino]methyl)phenyl)-6- (trifluoromethyl)pyrimidin-4(3H)-one |
| 2-(1-[1-(3,5,5-trimethylhexyl)piperidin-4-yl]-1H-1,2,3-triazol-4-yl)ethanol |
| 2-(1-[1-(cycloheptylacetyl)piperidin-4-yl]-1H-1,2,3-triazol-4-yl)pyridine |
| 2-(3-[(4-acetyl-1,4-diazepan-1-yl)methyl]phenyl)-3,5,6,7-tetrahydro-4H- cyclopenta[d]pyrimidin-4-one |
| 2-[2-(azepan-1-ylmethyl)phenyl]-6-isobutylpyrimidin-4(3H)-one |
| SU 5402 |
| RU 24969 |
| Flupirtine maleate |
| NSC-95397 |
| 5-methyl-2-[3-(morpholin-4-ylmethyl)phenyl]-6-(trifluoromethyl)pyrimidin-4(3H)-one |
| Benzo[a]phenanthridine-10,11-diol, 5,6,6a,7,8,12b-hexahydro-, trans- [CAS] |
| 4-Amino-1,8-naphthalimide |
| 5,6-dimethyl-2-(3-[(4-phenylpiperazin-1-yl)methyl]phenyl)pyrimidin-4(3H)-one |
| 6-butyl-2-(2-[(2-methylpyrrolidin-1-yl)methyl]phenyl)pyrimidin-4(3H)-one |
| 5-methyl-2-[3-(thiomorpholin-4-ylmethyl)phenyl]-6-(trifluoromethyl)pyrimidin-4(3H)-one |
| 2-(3-[(4-acetylpiperazin-1-yl)methyl]phenyl)-5-methyl-6-(trifluoromethyl)pyrimidin-4(3H)- one |
| Irinotecan HCl trihydrate |
| L-694,247 |
| Manoalide |
| Doxorubicin |

**Supplementary Table 2**

| **Morpholinos (MO)** | |
|---|---|
| Name | Sequence (5' to 3') |
| Control_MO | CCTCTT ACCTCAGTTACAATTTATA |
| *elf1*_ MO_*1* | GTGACTGTATTGTACCTGATTGATT |
| *elf1*_ MO_*2* | TTGTGATTGAACTCACCAGCCCCAC |
| *elf2a*_ MO | TATAAGTGTGGACTTTACCTGCTGT |
| *elf2b*_ MO | CACTTTACACACACAAGCTGCTCAC |
| *elf3*_ MO | ACAACAGTTACTCACCATATCTCGC |
| **sgRNA** | |
| Name | Sequence (5' to 3') |
| elf2b_sgRNA | UGGCUAAAGUCGAGGGUCAG |
| **Genotyping Primers** | |
| Name | Sequence (5' to 3') |
| elf2b_F | ctatgaaaccatgggtagagc |
| elf2b_R | ctcgtaggtcttctctccta |
